# Structure-based similarity network accelerates the discovery of lysins as oral microbiome modulators targeting periodontal pathogens

**DOI:** 10.64898/2026.04.11.717860

**Authors:** Fangfang Yao, Jiajun He, Raphael Nyaruaba, Fangyuan Chen, JianNan Zhou, Hang Yang, Hongping Wei, Yuhong Li

## Abstract

Microorganisms significantly influence human health, and dysbiosis of the oral microbiome plays a critical role in the development and progression of both oral and systemic diseases. This highlights the urgent need for novel therapeutics targeting specific pathogens. Here, we presented a structure-based pipeline to efficiently identify potential phage-derived periodontal lysins (LysPds) from nearly one million proteins. We predicted the structures of candidate lysins using AlphaFold2 and developed an innovative structure-based similarity network to classify them into distinct clusters, each with unique functional properties. A systematic characterization of 16 representative LysPds from 11 superfamilies revealed that over 90% demonstrated potent antibacterial activity against key periodontal pathogens. Among these, LysPd078 was identified as a promising preclinical drug candidate, effectively reconfiguring microbiome communities while demonstrating significant efficacy and safety in mouse models of periodontitis and calvarial infection. Our findings highlight the effectiveness of structure-based similarity networks in exploring vast protein spaces and underscore the potential of LysPd078 as a targeted modulating agent for the oral microbiome.

## Introduction

Microbial communities have occupied and shaped our habitats and their organisms for over 3.5 billion years^1^. As the second-largest microbial community in the human body, the oral microbiome co-evolves with the host^2^. Its bidirectional interaction with the host plays a crucial role in host health^3,4^. Subgingival (plaque) microbiota in advanced periodontitis is dominated by Gram-negative bacteria, including *Porphyromonas gingivalis*, *Fusobacterium nucleatum*, *Treponema denticola*, *Tannerella forsythia*, *Prevotella intermedia*, and *Aggregatibacter actinomycetemcomitans*, triggering irreversible destruction of the supporting tissues surrounding the tooth^5–7^. Currently, periodontitis is a global public health concern with a prevalence as high as 50%, and severe periodontitis affects over 1.08 billion adults worldwide, imposing a substantial health and economic burden^8,9^. Furthermore, studies have showed that periodontitis pathogens can spread through bloodstream, oral-pharyngeal inhalation or immune evasion, affecting distant organs and leading to various systemic inflammatory diseases, including cardiovascular disease, rheumatoid arthritis and even cancers^10–12^. Therefore, it is important to find effective intervention measures to reduce the periodontitis pathogens and reshape the oral microbial community, thereby promoting oral health and systemic health.

Recently, phages and phage-derived lysins have attracted increasing attention due to their potent antimicrobial activity, prompting their advancement into the clinical development pipeline^13,14^. However, the isolation of phages is hindered by the challenge of fastidious and anaerobic culture requirements of most periodontitis pathogens. To date, only a few phage strains targeting *Aggregatibacter* and *Fusobacteria* species have been isolated^15^. In contrast, phage-derived lysins function as peptidoglycan hydrolases, acting differently from phages and presenting lower risks of resistance and biosafety concerns^16^. Additionally, lysins as protein drugs have a well-established regulatory pathway for drug development^17^. While significant progress has been made in clinical research on lysins targeting pathogenic bacteria^18,19^, no lysins targeting periodontitis pathogens have been reported.

Traditional lysin discovery relies on isolating and sequencing phages, a time-consuming and labor-intensive process with unpredictable outcomes. Bacterial genomes contain numerous molecular imprints left by phages, known as prophages, embodying a vast and unexplored potential reservoir of lysins^20,21^. Despite the abundance of data in databases, the complexity and low-throughput nature of bio-experiments necessitate accurately identifying representative sequences to reduce workload and maximize the exploration of functional space. However, billions of years of natural selection have shaped the functional diversity and hybridization of protein sequences^22^. Additionally, frequent recombination events throughout phage evolution have rendered lysins increasingly hybridized^23^, making their identification more challenging. In fact, three-dimensional structures provide a more precise mapping of protein function compared to one-dimensional sequences. Recent advancements in artificial intelligence have led to unprecedented accuracy in predicting protein structures from amino acid sequences^24–26^, providing the possibility for large-scale exploration of protein function based on structure. However, current structure-based clustering methods tend to introduce bias when dealing with unevenly distributed samples^27,28^.

In this study, we conducted the first large-scale exploration of potential lysins within nearly a million proteins from periodontitis pathogens. We proposed the structure-based similarity network to effectively explore protein functional spaces with hybrid and unevenly distributed features. Using this approach, we identified novel periodontal lysins (LysPds) that demonstrated robust antibacterial activity against six major periodontitis pathogens. Among these, LysPd078 targeted periodontitis pathogens and promoted the microbiota of periodontitis subgingival plaque toward a healthier state. Moreover, LysPd078 demonstrated remarkable therapeutic efficacy and safety in mouse models of osteolysis induced by bacterial infection. These findings highlight the immense potential of structure-based similarity networks in discovering and decoding proteins with novel functions, providing valuable insights for future explorations and innovations in drug development.

## Result

### Discovery and classification of novel LysPds based on structure

The study of periodontitis pathogens has spanned decades. Despite extensive efforts in this field, phages isolated from periodontitis pathogens are exceedingly rare, raising questions about their prevalence^15^. To explore potential phage-derived lysins in the genomes of these pathogens, we employed advanced machine learning tools to investigated the presence of prophages in these bacteria **(Fig. 1a)**. We found that over half of the periodontitis pathogens (191/364) contained recognisable integrated phages, with some strains even holding up to 10 prophages (**Fig. 1b-d**). This finding highlights the feasibility of directly mining phage-derived lysins from the genomes of periodontitis pathogens. However, lysins are a large and highly diverse group of enzymes characterized by functional diversity and extensive hybridity^23^, making it difficult to classify by sequence alone. To overcome this, we developed a structure-based workflow to efficiently discern representative lysins **(Fig. 1e and Extended Data Fig. 1a)**. Using a set of seed lysins (**Supplementary Data Table 2**), we screened nearly one million proteins from periodontitis pathogens. Following manual curation, we used AlphaFold2 to predict three-dimensional structures and ultimately obtained 164 LysPds with reliable structures.

**Fig. 1:**
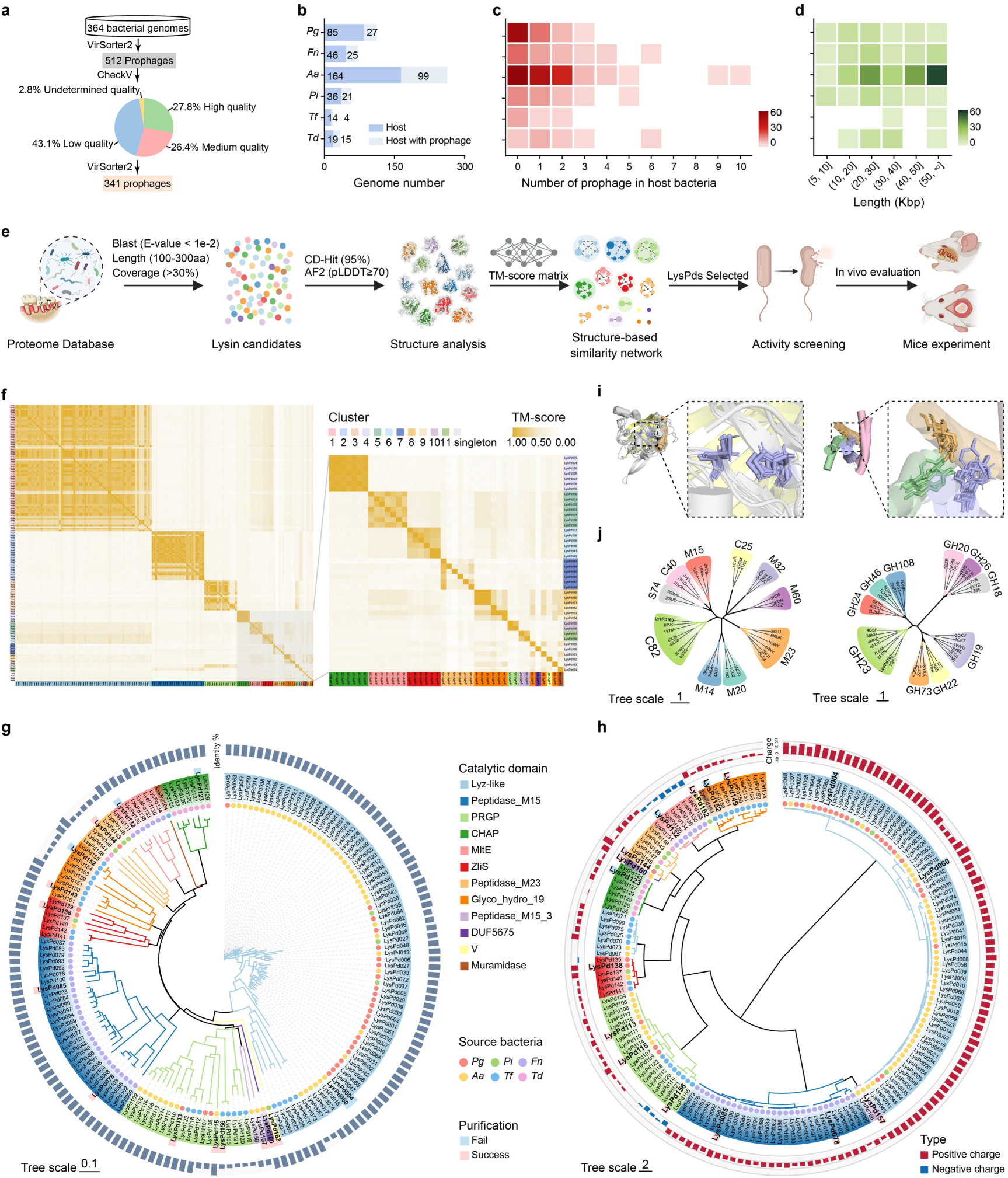
Protein classification of LysPds based on structures. a, Workflow for identifying prophages (length ≥5 kbp) against periodontitis-associated bacteria. b, The number of host genomes that containing prophages. c, Distribution of prophage numbers in different host genomes. d, Length distribution of prophage genome in different host. e, Workflow for mining LysPds against periodontitis-associated bacteria. f, Structural similarity matrix reflecting similarities among the 164 LysPds structures. The orange-to-white heatmap indicates the degree of structural similarity. Cluster numbers are distinguished by different colors and are marked along the vertical sides of the matrix. At the bottom, colors denote different types of catalytic domains in the corresponding LysPds. g, Sequence-based dendrogram of the 164 LysPds, with different colors representing each catalytic domain. The length of the external outer dark blue bars represents the identity between the LysPd and the recognizable oral microbiome phage protein. h, Structure-based dendrogram of the 164 LysPds. Outer bar lengths represent the net charge of LysPds, with positive charges indicated in red and negative charges in blue. i, Structural alignment of LysPd160 (DUF5675) with typical C82 peptidases (left) and LysPd162 (V) with typical GH23 glycosidases (right). Conserved catalytic residues displayed as stick models, with additional protein domains removed for clarity. j, Structure-based dendrogram of LysPd160 with typical peptidases (left) and LysPd162 with typical glycosidases (right).

Next, we evaluated the structure similarity of 164 candidate LysPds via US-align. A structure similarity matrix was generated based on all-against-all alignment (Fig. 1f). We conduct a structural dendrogram that classified 164 candidate lysins into distinct branches, each characterised by unique structures corresponding to protein functional attributes **(Fig. 1h)**. Notably, the structure-based dendrogram outperformed its sequence-based counterpart in decoding enzyme functions, correctly grouping Peptidase_M15_3 lysins and Muramidase lysins, which had been misclassified by sequence alone **(Fig. 1h, g).** Moreover, LysPd160 (DUF5675 domain) and LysPd162 (V domain) formed isolated branches in the sequence dendrogram, but were clustered with peptidases and glycosidases in the structural dendrogram. Although these domains are widely distributed across phages and over 500 bacterial genera, their functions still unknown **(Extended Data Fig. 2a-d)**. Further structural analysis revealed that these two domains share some similarities with their respective typical GH23 and C82 families but exhibit significant differences **(Fig. 1i, j and Extended Data Fig. 3)**. We therefore propose classifying them as GH23-like and C82-like families. This clearly demonstrated that AI-assisted structure prediction offered more robust solution for decoding protein functions compared to sequences.

### Clustering of LyPds using the structure-based similarity network

Due to natural distribution and sequencing limitations, biological data frequently exhibit a severe imbalance between diversity and quantity^22^. Conventional clustering methods tend introduce random biases, leading to discrepancies between clustering results and actual functional groupings **(Fig. 2a)**. To address this, we developed a new clustering method: structure-based similarity network **(Fig. 2b)**. In this network, edges are formed between nodes (representing proteins) when their structural similarity exceeds a predefined threshold, enabling objective clustering of proteins with similar structures. This method provides an accurate mapping between protein functions and clustering while maintaining consistent differentiation between distinct clusters. Using this approach, the 164 candidate LysPds were clustered into 11 clusters and 6 singletons **(Fig. 2c)**. Notably, certain lysin families with functional differences were correctly divided into separate clusters, such as M15 lysins and M15_3 lysins **(Extended Data Fig. 1b)**. Both PGRP lysins and GH19 lysins were divided into different clusters with distinct structural features, suggesting potential functional properties variations within each group **(Extended Data Fig. 1b)**. Given the importance of charge in the activity of lysins against Gram-negative bacteria, we selected highly charged LysPds from each cluster for further study **(Fig. 2d, e)**. These LysPds demonstrated uniform diversity and significant novelty, with as low as 25% identity to known lysins, greatly expanding the known range of lysins **(Extended Data Fig. 1c-e and Supplementary Data Table 3)**.

**Fig. 2:**
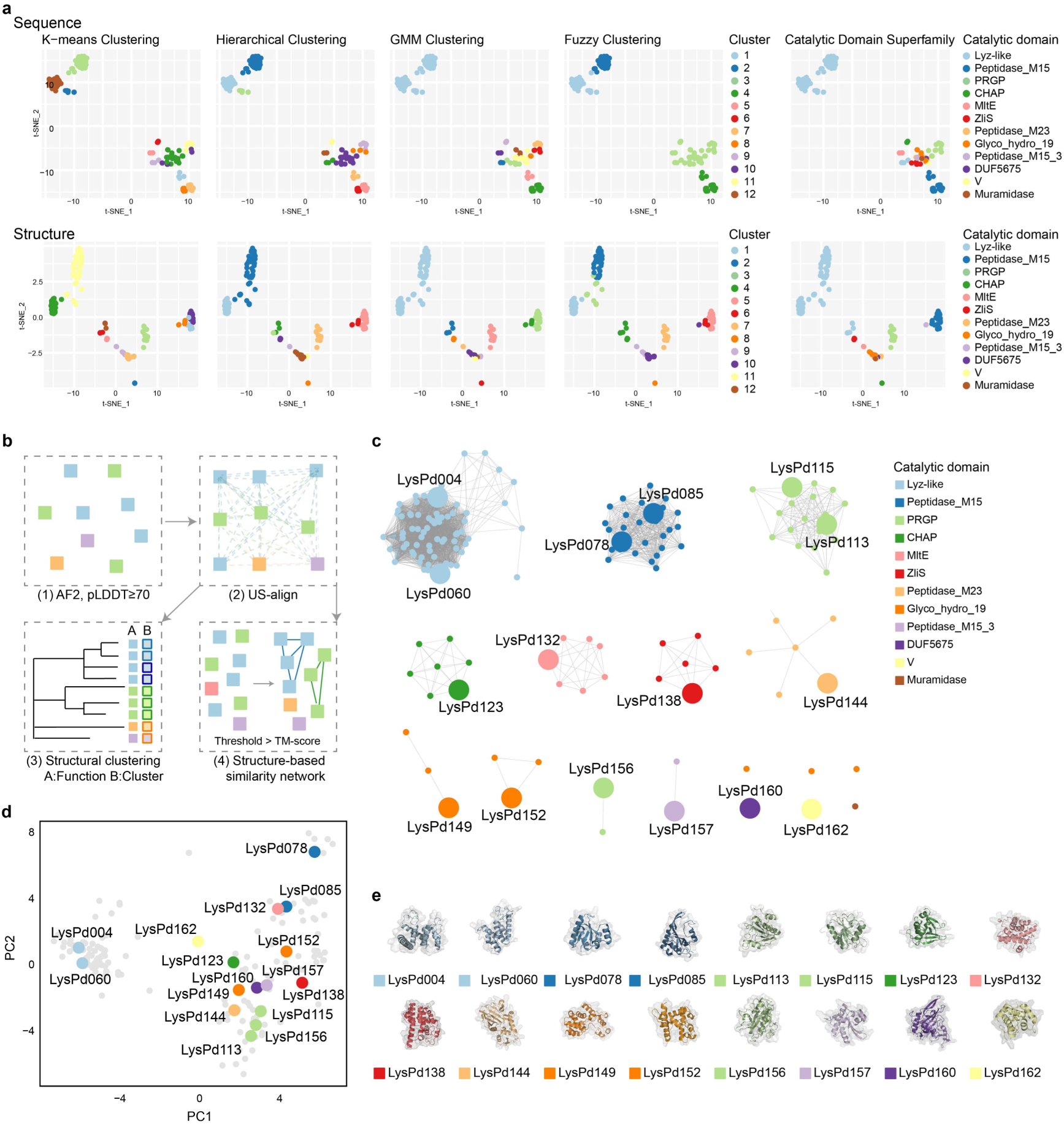
Structural clustering of newly discovered LysPds. a, Clustering results of LysPds based on sequence (Upper panels) and structure (Lower panels) using various methods. The results were colored according to different clustering results or catalytic domain. The default maximum number of cluster components was set to 12. b, Schematic diagram of the structure-based similarity network. c, The structure-based similarity network represented LysPds as nodes (circles) and all pairwise structure relationships (alignments) that exceed the threshold (TM-score 0.75) as edges (lines). The 164 LysPds were divided into 11 clusters and 6 singletons, coloured based on catalytic domains. Selected LysPds are enlarged and labeled with their respective names. d, PCA of the 100-dimensional eigenvectors of the 164 LysPds. e, Predicted structures of 16 selected LysPds.

### Screening potent LysPds against periodontitis bacteria

The 16 LysPds identified from our clustering pipeline were tested for antibacterial activity against six key periodontal pathogens (i.e. *P. gingivalis*, *F. nucleatum*, *T. denticola*, *T. forsythia*, *P. intermedia*, and *A. actinomycetemcomitans*). Following multiple attempts, 11 lysins were successfully expressed as soluble proteins in *E. coli* **(Extended Data Fig. 4a)**. Among these, 10 lysins exhibited highly antimicrobial activity, resulting in a notable success rate of 90% **(Fig. 3a)**, with correlating linearly with their net charges **(Extended Data Fig. 4b)**. Utilizing *P. gingivalis* and *F. nucleatum* as model bacteria, the impact of LysPds on the bacterial cell wall, cytoplasmic membrane, and outer membrane was evaluated, revealing variances in outer membrane permeability that paralleled their antimicrobial activity **(Fig. 3b and Extended Data Fig. 4c)**. Following the initial screening, the top four LysPds (LysPd078, LysPd138, LysPd144 and LysPd157) with the highest activity were further analyzed. LysPd144 reduced *F. nucleatum* by 99.97% within 5 minutes **(Fig. 3c, Time)**, while the MBC of LysPd078 against *P. gingivalis* and LysPd144 against Fusobacterium nucleatum was 25 μ g/mL (0.78-1.77 μ M) **(Fig. 3c, Concentration and Extended Data Fig. 4d)**. Remarkably, these four LysPds retained their antimicrobial activity after incubation at temperatures up to 100°C **(Fig. 3c, Temperature)**, demonstrating thermotolerance exceeding that of lysins from thermophilic phages **(Extended Data Table 1)**. Additionally, all of these thermostable enzymes exhibited strong storage stability, retaining their activity after long-term storage (**Extended Data Fig. 5a)**. Specifically, LysPd078 preserved nearly 99% of its activity after 6 months of storage at 22°C and 37°C **(Fig. 3d)**.

**Fig. 3:**
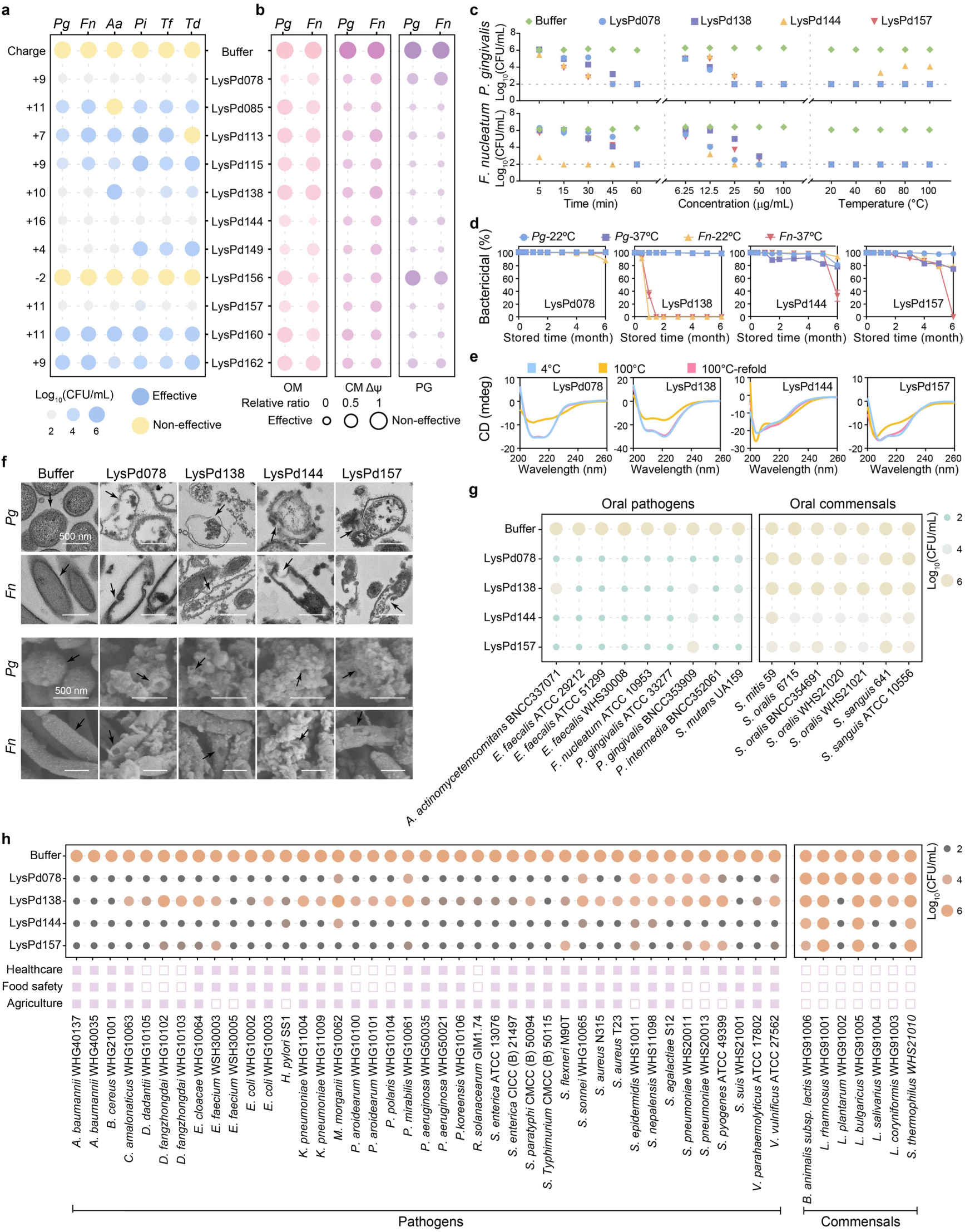
Antibacterial activity of selected LysPds. a, Antibacterial activities of LysPds against six bacterial strains (*P. gingivalis* W83, *F. nucleatum* ATCC 25586, *T. denticola* ATCC 35405, *T. forsythia* ATCC 43037, *P. intermedia* ATCC 25611, and *A. actinomycetemcomitans* HK1651) used for the initial screening. The LysPds were highly effective (P ≤ 0.0001 in Dunnett’s test, blue) against periodontitis pathogens. b, Effect of LysPds on the bacterial cell wall (PG), cytoplasmic membrane (CM Δψ), and outer membrane (OM) under various conditions. c, The effect of time, concentration, and temperature on the bactericidal activity of LysPds. d, Storage stability of LysPds at 22°C and 37°C. e, Circular dichroism spectra of LysPds at 4°C, 100°C, and 100°C-refold (heating at 100°C for 1h followed by cooling to 4°C). f, TEM (Upper panels) and SEM (Lower panels) images showed the effects of LysPds on the bacteria scale bar, 500 nm. g, Antimicrobial activity of LysPds against various strains of oral pathogenic bacteria and oral commensal bacteria. h, Antimicrobial activity of LysPds against various strains of different industries. Data are presented as mean values ± SD.

In order to elucidate the thermostability mechanism of the novel LysPds, circular dichroism (CD) was utilized to monitor changes in their secondary structure at various temperatures and determine their melting temperature (Tm). CD analysis revealed that although the denaturation temperatures of LysPds were below 100°C, the secondary structures almost completely recovered after rapid cooling from 100°C **(Fig. 3e and Extended Data Fig. 5b, c)**, indicating that their thermostability was associated with reversible thermal denaturation. Notably, the thermophile scores of proteins derived from periodontal-related bacteria were significantly higher than those from other pathogens **(Extended Data Fig. 5d, e and Extended Data Table 2)**, suggesting the heat stability of these proteins may not incidental.

Subsequently, to further explore the antimicrobial effect of LysPds on bacteria, we employed transmission electron microscopy (TEM) and scanning electron microscopy (SEM) to observe the changes in bacterial microstructure. After enzyme treatment, the integrity of the bacterial cell wall was destroyed, with collapsed and wrinkled cell borders, showing that the enzymes directly induced lysis of bacteria **(Fig. 3f)**. Then, we tested the antibacterial spectrum of LysPds to assess their potential role in regulating oral microbiota balance. Interestingly, while LysPd078 showed minimal activity against oral commensals, it exhibited potent antimicrobial activity against other oral pathogens **(Fig. 3g)**. Moreover, LysPds demonstrates broad-spectrum bactericidal effects against more than 30 species that pose major challenges in the healthcare, food, and agriculture industries, while exhibiting minimal activity against recognized probiotics **(Fig. 3h).** These results demonstrated that the identified LysPds, particularly LysPd078 and LysPd144, exhibited exceptional antimicrobial efficacy and application potential.

### The efficacy of LysPd078 and LysPd144 against biofilm and subgingival plaque

In periodontitis, pathogens primarily exist as biofilms rather than planktonic forms, and the pattern of co-aggregation of *P. gingivalis* and *F. nucleatum* mutually promotes their survival and proliferation^29,30^. This plaque-biofilm forms a barrier, greatly impeding the penetration of antimicrobial drugs. To assess the efficacy of LysPds in targeting biofilms, LysPd078 and LysPd144 were applied to target single- and dual-species biofilm models of *P. gingivalis* and *F. nucleatum*. Results from CFU counting, SEM, and CLSM analysis demonstrated that these LysPds reduced up to 99.9% of live bacteria within the biofilms, with most cells appearing incomplete, distorted, and wrinkled, exhibiting red fluorescence indicative of cell death **(Fig. 4a-c and Extended Data Fig. 6a-c)**.

**Fig. 4:**
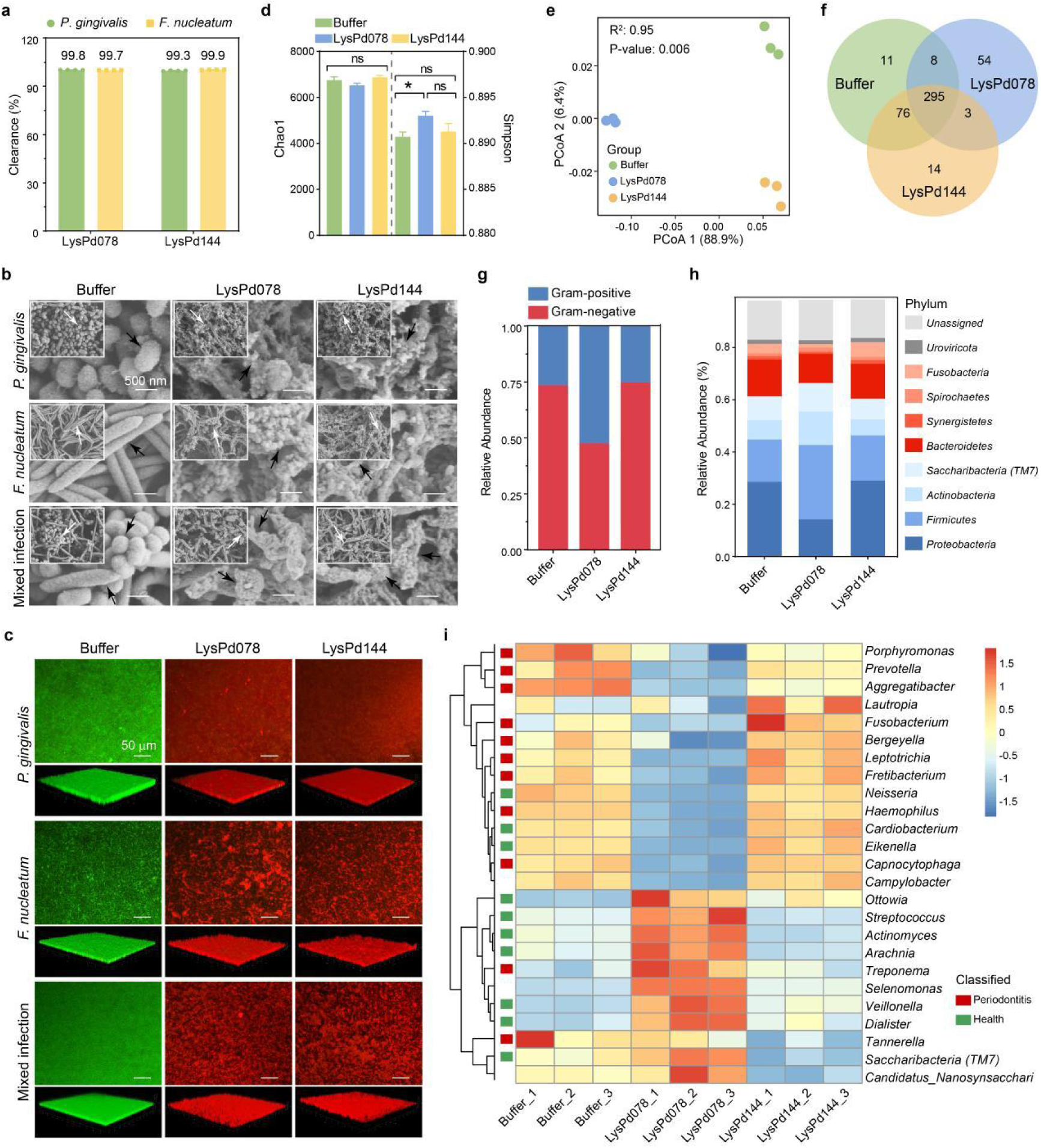
Bactericidal effects of LysPd078 and LysPd144 on biofilm models and subgingival plaque. a, Change in bacterial load measured as percentage intra-biofilm bacteria killed by LysPd078 and LysPd144 respectively. b, Morphological changes in single- and dual-species biofilms treated with LysPd078 and LysPd144 and visualized by SEM. Scale bar, 500 nm. c, Representative images of live/dead fluorescent staining of biofilms treated with LysPd078 and LysPd144, where green fluorescence indicates live bacteria, and red fluorescence indicates dead bacteria. Scale bar, 50 μm. d, Richness and diversity in the microbiome of subgingival plaque across different treatment groups. e, Principal co-ordinate analysis (PCoA) illustrating differences in microbiome compositions among groups. f, Venn diagram showing the overlap of subgingival flora among the different groups. g, Distribution of species-level taxa by Gram staining characteristics across the different groups. h, Stacked bar plot depicting the mean relative abundance of the top 10 taxa at the phylum level, with periodontitis-associated taxa in red and health-associated taxa in blue. i, Heatmap generated by hierarchical clustering of the differentially abundant genera. The color gradient from red to blue indicates the degree of normalized relative abundances. Genera significantly associated with periodontitis are marked in red, while those associated with health are marked in green. Statistical significance was calculated via Permutation test; *p < 0.05, ns, not significant. Data are presented as mean values ± SD.

Further, the antimicrobial potential of LysPd078 and LysPd144 was explored in clinical subgingival plaque samples from real-world evidence. These treated samples from periodontitis patients were further incubated propidium monoazide (PMAxx), which was used to distinguish live and dead bacteria, thus enabling the analysis of changes in the oral microbiota. We found that treatment with LysPd078 promoted the simpson diversity index of the plaque-biofilm (**Fig. 4d**), accompanied by significant dissimilarity in microbiome composition (**Fig. 4d, f**), suggesting its potential to alter the microbial composition. Notably, LysPd078 significantly decreased the abundance of Gram-negative bacteria (**Fig. 4g**), which are closely associated with periodontitis. And LysPd078 promoted a shift in the microbiota towards a healthier state^31,32^ by increasing the relative abundance of health-associated taxa and reducing disease-associated ones at both the phylum and genus levels (**Fig. 4h, i**). At the species level, the abundance of some key pathogenic bacteria also decreased, while the proportion of commensal bacteria increased (**Extended Data Fig. 6d, e**). This shift caused by LysPd078 lowered the pathogenicity of the oral microbiome and transitioned the microbial composition toward a healthier state.

### LysPd078 and LysPd144 alleviate osteolysis induced by bacterial infection

Following the satisfactory antimicrobial activity exhibited by LysPd078 and LysPd144, their in vivo efficacy was further investigated in the mouse periodontitis model (**Fig. 5a**) and the calvarial infection model, both induced by *P. gingivalis* and *F. nucleatum* (**Fig. 5f**). Cytotoxicity assays conducted before in vivo testing indicated that LysPd078 and LysPd144 were cytocompatible at concentrations of up to 1000 μ g/mL.(**Extended Data Fig. 7a**).

**Fig. 5:**
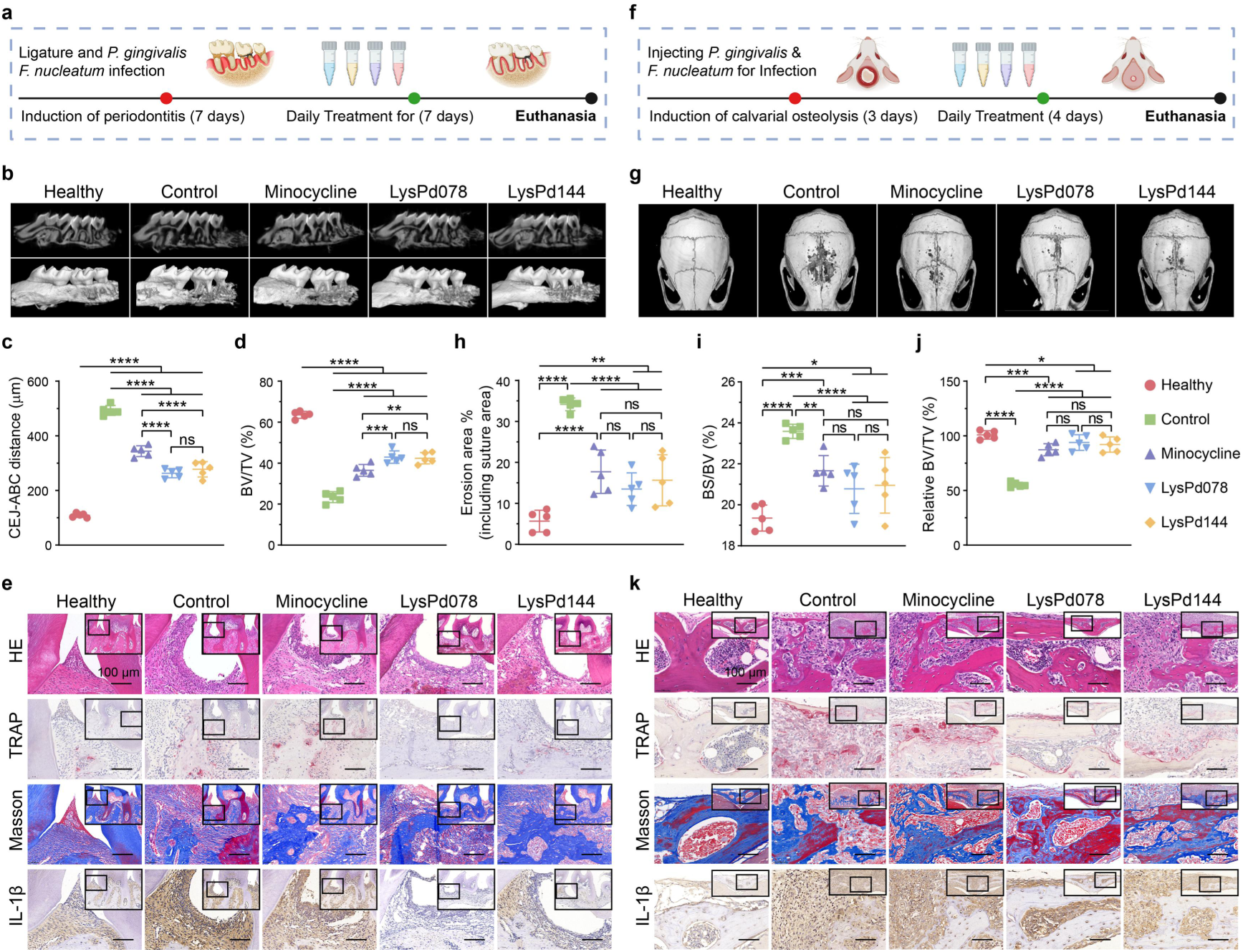
In vivo alleviation of bacterial-induced osteolysis by LysPd078 and LysPd144. Schematic illustration of the periodontitis model (a) and calvarial infection model (f), depicting the experimental design and treatment process. b, Micro-CT reconstructed images of the right maxillary molar area from the indicated groups of mice. Both multiplanar reconstruction (Upper panels) and three-dimensional volume (Lower panels) are shown. c, Quantification of the distance from the cementoenamel junction (CEJ) to the alveolar bone crest (ABC). d, Ratio of bone volume (BV) to total volume (TV) of in the maxilla surrounding the second molar. e, Histology staining images of periodontal tissue showing the pathogenic alteration. Scale bar, 100 μm. g, Three-dimensional micro-CT reconstructed images of calvarial bone. h, Quantification of the erosion area in the calvaria, including the suture calvarial area centred on the intersection of the coronal and sagittal sutures. i, Ratio of bone surface (BS) to bone volume (BV) of calvaria surrounding the erosion area. j, Relative ratio of bone volume (BV) to tissue volume (TV) of the calvaria compared to the healthy group. k, Representative histology staining images of calvarial bone showing the erosion area. Scale bar, 100 μm. Statistical significance was calculated according to One Way ANOVA; n = 5/group, *p < 0.05, **p < 0.01, ***p < 0.001, ****p < 0.0001, ns, not significant. Each data point represents a biologically independent mouse.

In the mouse periodontitis model, untreated mice showed significant alveolar bone loss. Notably, LysPd078 demonstrated efficacy comparable to LysPd144 in reducing bone loss, but superior to minocycline, the most widely used antibiotic for periodontitis treatment (**Fig. 5b-d**). Histopathological analysis revealed significant loss of epithelial attachment and infiltration of inflammatory cells in the periodontal tissues of the control groups. In contrast, both LysPd078 and LysPd144 alleviated histological inflammatory manifestations, more effectively than Minocycline. As evidenced by fewer positive stains in tartrate-resistant acid phosphatase (TRAP) staining, and an increase in collagen fibers, as seen in Masson’s trichrome staining (**Fig. 5e**). Additionally, treatment with LysPd078 and LysPd144 significantly reduced levels of the proinflammatory cytokine IL-1β compared to the control group. Importantly, no organ damage or differences in body weight were observed following local treatment with either LysPd078 or LysPd144 (**Extended Data Fig. 7b, c**), demonstrating their potential for safe and effective periodontitis treatment.

In the mouse calvarial infection osteolysis model, untreated mice exhibited significant increase in calvarial bone resorption. Treatment with LysPd078, LysPd144, and minocycline showed comparable efficacy in reducing bacteria-induced bone loss (**Fig. 5g-j**). This reduction in bone loss was associated with inhibited inflammation and suppression of osteoclast formation on the calvarial bone surface, consistent with results from the periodontitis model (**Fig. 5k**). These findings highlight LysPd078 and LysPd144 as promising candidates for treating periodontitis and related bone loss conditions.

## Discussion

Microorganisms form the backbone of ecosystems, playing critical roles from the smallest bacteria to complex human systems. The periodontal environment, in particular, is a highly intricate and densely populated ecosystem, hosting billions of bacteria from over 700 species^33^. Maintaining a balanced oral microbiome is essential for both oral and systemic health. However, when this balance is disrupted by pathogenic bacteria, it can lead to a range of diseases, from localized periodontal issues to broader systemic conditions such as cardiovascular disease and diabetes^10,12^. Traditional antibacterial methods often struggle with limited spectra and can inadvertently disrupt microbial balance, highlighting the urgent need for novel approaches to restore and maintain periodontal health^10^. Phage-derived lysins offer a promising solution due to their potent bactericidal activity and diverse-spectrum effectiveness^16^. In this study, we leveraged advanced computational tools and big data to overcome the challenges of isolating phages from fastidious anaerobic bacteria. By analyzing nearly one million proteins from periodontitis pathogens, we identified a comprehensive set of lysins targeting key periodontal bacteria.

Facing the exponential growth of genomic data, effectively and comprehensively exploring the entire functional space poses a tremendous challenge. The highly diverse enzymes in nature exhibit significant sequence hybridity due to frequent gene recombination during their evolution. This makes it extremely challenging to delineate protein function based solely on sequence. In contrast, the three-dimensional structure of proteins provides valuable functional insights and a more stable reference framework for understanding their roles^34^. In recent years, significant advancements in artificial intelligence have revolutionized high-precision protein structure prediction, inspiring us to delve deeper into the protein functional space. Using a new structure-based clustering method, we effectively categorized lysins into distinct functional groups, and even decoded lysins with previously unknown functions. This approach demonstrates the superior capability of structural analysis in overcoming the limitations of sequence-based methods, particularly in accurately classifying proteins and decoding their unknown functions.

In fact, recent studies underscore the huge potential of structure-based clustering in exploring protein functions^27,28,35^. Similar to these studies, we found that mainstream clustering methods could efficiently cluster sequence-heterogeneity proteins when dealing with carefully selected uniform data (**Extended Data Fig. 2e, f**). However, practical protein exploration often involves large, uneven datasets from nature^22^. Those clustering methods tend to overly focus on similar predominant proteins, often regarding novel, rare functional sequences as noise or outliers. This can lead to erroneous divisions that fail to provide reasonable partitions while ensuring accurate functional mapping. In contrast, by setting objective thresholds, the structure-based similarity network offers a robust solution for uniform division and accurate clustering within diverse and uneven datasets. Particularly, once the mapping relationship between structure and function is established, this approach shows unprecedented advantages due to the shared highly similar structural skeletons within clusters.

Among the 16 LysPds selected from different clusters using the structure-based similarity network, 10 of the 11 successfully expressed lysins displayed strong antibacterial activity against periodontitis pathogens and exhibited distinct functional properties. Notably, four of these LysPds demonstrated exceptional thermostability, a critical feature for protein therapeutics, particularly in impoverished but healthcare-burdened regions like Africa and Latin America where cold chain logistics are challenging^36^. Moreover, the antibacterial spectrum of the four LysPds not only includes oral pathogens but also covers most bacteria listed on the 2024 WHO Bacterial Priority Pathogens List^37^. And LysPds also demonstrated a potent ability to eradicate notorious pathogens in the agricultural and food industries, such as R. solanacearum^38^ that causes a 20-100% crop yield losses worldwide and *Salmonella spp.*^39^ that results over 90 million foodborne cases each year. These results highlight the broad potential of LysPds in oral health and various industries.

Periodontitis is a chronic, polymicrobial disease that requires targeted intervention to restore microbial balance. When evaluating the antibacterial spectra of LysPds, we found that LysP078 displayed highly specific and efficient antibacterial activity against oral pathogens, suggesting its potential to modulate the oral microbiota. Metagenomic sequencing of plaque samples from periodontitis patients further revealed that LysPd078 selectively targeted and reduced pathogenic bacteria in subgingival plaque, facilitating a shift towards a healthier oral microbial community. Furthermore, LysPd078 exhibited excellent biocompatibility and effectively inhibited bone destruction in vivo, suggesting their potential as safe and effective treatments for periodontitis. Given the close association between periodontal microbiota and systemic health, we propose that LysPd078 not only holds promise as a drug candidate for treating periodontitis but may also reduce related systemic health risks. Since the elucidation of the DNA double helix model in 1953, decades of research utilizing a limited subset of cultivable microorganisms have fueled the current thriving multi-trillion-dollar biotechnology market^40^. Today, faster and cheaper genome sequencing technologies are accelerating us into a new era of planetary-scale genomics, encompassing over 8 billion non-redundant proteins^34^. Faced with such an extensive sequence space, there is an urgent need to efficiently identify representative proteins to address the conflict between experimental throughput and data. Fortunately, we are now witnessing a scientific research paradigm transformation empowered by AI for science^41^, encouraging us to discover new drugs^42–44^, materials^45^, and even extraterrestrial intelligence^46^ with unprecedented efficiency.

Here, with AI-assisted structural clustering, we explored the nearly one million proteins of periodontal bacteria. And experimental characterization of only 16 proteins led us to discover the preclinical drug candidate LysPd078, which holds potential for reshaping the periodontal microbiome. This fully demonstrates the power of structure-based similarity network to achieve rational functional clustering in diverse and uneven datasets. More importantly, as structural prediction accuracy continues to improve, structural clustering at the residue level of specific protein positions will provide possibilities for drawing omics-scale protein-protein and protein-drug interaction networks^47–49^, further deepening our understanding of life^50^. Overall, we believe that structure-based clustering will fundamentally transform the discovery of functional proteins, ushering us into the era of AI-driven scientific revolution.

## Methods

### Bacterial strains and culture conditions

Bacterial strains, their cultivation media, growth conditions, and source are listed in Supplementary Table S1. Bacteria were cultivated under aerobic, microaerobic (5% CO_2_), and anaerobic conditions (AnaeroPack, Mitsubishi) as required.

### Identification of prophages

A total of 364 genomes from key periodontitis pathogens, including *Porphyromonas gingivalis*, *Fusobacterium nucleatum*, *Treponema denticola*, *Tannerella forsythia*, *Prevotella intermedia*, and *Aggregatibacter actinomycetemcomitans* were downloaded from NCBI as of January 2022. To identify potential prophages within these genomes, we followed a standardized protocol (DOI dx.doi.org/10.17504/protocols.io.bwm5pc86). Here, potential prophages in the genomes were identified using VirSorter2^51^and CheckV^52^. Additionally, prophage proteins identified using geNomad^53^ were compared with proteins from the Oral Virus Database (OVD)^54^ as part of the recognizable oral microbiome phage protein.

### Mining and Clustering of Lysins

The proteins of major periodontitis pathogens were downloaded from NCBI as of January 2022. A carefully selected set of experimentally validated Gram-negative lysins (Supplementary Data Table 2) were used as queries for BLAST searches (E-value < 1e-2) against periodontitis pathogen proteins, filtering out hits with coverage below 30%. Considering the length distribution of Gram-negative lysins^55^, 796 hits ranging from 100 to 300 amino acids in length were retained as candidate lysins. The sequences were then de-redundant with 95% identity using CD-Hit^56^, followed by predicting their three-dimensional structures using AlphaFold2 and proteins with reliable structures (pLDDT ≥ 70) were further analysed.

All-against-all alignments of LysPds were performed using US-align^57^ to evaluate their structural similarity. TM-scores were extracted to construct the structural similarity matrix and generate the hierarchical clustering dendrogram, which was visualized using tvBOT^58^. For comparison, a sequence-based dendrogram was also generated (https://www.ebi.ac.uk/jdispatcher/msa/clustalo) and the sequence similarity matrix was calculated by TB-tools^59^. Batch CD-Search against the CDD^60^ database was conducted to identify the superfamilies of catalytic domains within the LysPds.

Based on the structural similarity matrix, the structure-based similarity network was generated using a TM-score threshold of 0.75 to divide LysPds into different clusters, visualized in Cytoscape. For comparison, other mainstream clustering methods (K-means, Hierarchical, Gaussian Mixture Model, Fuzzy) were also used to cluster LysPds. Furthermore, t-SNE dimensionality reduction was applied to visualize the structural and sequence similarity matrices, which were colored according to the different clustering outcomes or catalytic domain types.

Based on the structure-based similarity network, LysPds with the highest charge were selected from each cluster as representatives, and their structures were visualized using PyMOL. The selected LysPds were highlighted in the PCA plot, generated by dimensionality reduction of the 100-dimensional eigenvectors of all 164 LysPds^61^. To determine their novelty, the union set of selected LysPds and reported lysins was de-redundant based on different sequence similarity thresholds (https://web.expasy.org/decrease_redundancy/).

### Analysis of LysPd160 and LysPd162

Protein sequences and architectures containing DUF5675 (C82-like) and V (GH23-like) domains were downloaded from NCBI. As previously described, structural alignments were performed using US-align for LysPd160 with typical glycosidases and LysPd162 with typical peptidases, and structural dendrograms were generated from structural similarity matrices. t-SNE plots were generated using structural similarity matrix and sequence similarity matrices as previously described. These plots were colored according to different clustering results to evaluate their effectiveness in analyzing carefully selected uniform data.

### Plasmid construction and Protein purification

LysPds were commercially synthesized (Zhonghe Gene), and cloned into the NcoI and XhoI sites of pET-28b(+) with a 6×His tag at the C-terminus. The recombinant proteins were expressed in *E. coli* BL21 (DE3) as described previously^62^. Briefly, bacterial cells were lysed (high-pressure cell disruptor), centrifuged (12,000g; 30 min) and filtered (polyvinylidene difluoride, 0.22-μm pore size). Sample-loaded HisTrap FF columns were washed and eluted with 20 and 250 mM imidazole, respectively. After dialysis against 10 mM HEPES (pH 7.4), purified proteins were stored at 4°C until use. Protein quantification and 15% sodium dodecyl sulfate-polyacrylamide gel electrophoresis (SDS-PAGE) were performed following standard protocols^62^.

### Outer membrane permeabilization assays

The membrane permeability of LysPds was determined by using the N-phenyl-1-naphthylamine (NPN) uptake assay^63^. Briefly, bacteria were washed and resuspended at 10^8^ colony-forming unit (CFU)/mL in buffer (5 mM HEPES, 5 mM glucose, pH 7.4), and then incubated with 100 μg/mL LysPds in the presence of 10 μM NPN in a white 96-well plate at 37°C for 15 min. The fluorescence intensity was measured using a Synergy H1 microplate reader (BioTek, Winooski, VT) set at 350 nm for excitation and 420 nm for emission. Bacterial cells without NPN (buffer) were used as a negative control for background fluorescence. The following equation was used to reflect the ratio of the difference between buffer and LysPds: Ratio (relative permeable outer membrane activity) = (FI_sample_-FI_nc_)/(FI_buffer_-FI_nc_), where FI_nc_ = fluorescence of the negative control, FI_buffer_ = fluorescence of the buffer treated, and FI_sample_ = fluorescence of the LysPds treated. All experiments were performed in triplicate.

### Cytoplasmic membrane depolarization assay

The depolarization activity of LysPds on the cytoplasmic membrane was assessed through the uptake of the cationic membrane potential-sensitive dye DiSC_3_(5)^64^. Briefly, bacteria were grown overnight, harvested, washed, and re-suspended at 10^7^ CFU/mL in the same buffer (5 mM HEPES, 20 mM glucose, pH 7.4) but containing 100 mM KCl and 1 µM DiSC_3_(5). Bacteria were incubated for 30 min to allow stable uptake and quenching of the dye in the bacterial membrane. Samples were incubated with 100 μg/mL LysPds in a white 96-well plate at 37°C for 15 min. The fluorescence intensity was measured using Synergy H1 microplate reader at 620 nm for excitation and 670 nm for emission. Bacterial cells without DiSC_3_(5) were used as a negative control for background fluorescence. The following equation was used to reflect the ratio of the difference between buffer and LysPds: Ratio (relative cytoplasmic membrane depolarization activity) = (FI_sample_-FI_nc_)/(FI_buffer_-FI_nc_), where FI_nc_ = fluorescence of the negative control, FI_buffer_ = fluorescence of the buffer treated sample, and FI_sample_ = fluorescence of the LysPds treated sample. All experiments were performed in triplicate.

### Crude peptidoglycan purification and muralytic activity assay

Crude peptidoglycan was used to determine muralytic activity as previously described^65^. Biomass of *P. gingivalis* W83 and *F. nucleatum* ATCC 25586 was harvested from the stationary growth phase and boiled in 5% SDS to lyse the cells with vigorous shaking. After overnight incubation at room temperature, the suspension was washed and concentrated to an optical density (OD_600_) of ∼1.5 in MilliQ water. Crude peptidoglycan was then added to LysPds at a concentration of 100 μg/mL and incubated at 37°C for 1 h. The OD_600_ was measured using a Synergy H1 microplate reader, with 10 mM HEPES buffer without crude peptidoglycan serving as a negative control for background fluorescence^66^. The following equation was used to reflect the ratio of the difference between buffer and LysPds: Ratio (relative muralytic activity) = (OD_buffer_-OD_nc_)/(OD_sample_-OD_nc_), where OD_nc_ = OD_600_ of the negative control, OD_buffer_ =OD_600_ of the buffer treated sample, and OD_sample_ = OD_600_ of the LysPds treated sample. All experiments were performed in triplicate.

### Halo assay

Briefly, bacteria overlay agarose^67^ was prepared by culturing 100 mL of each bacterium overnight. After centrifugation (8,000 rpm, 5 min), the bacteria were suspended in 35 mL 10 mM HEPES with 0.7% agarose, autoclaved, poured over plates, and stored at 4 °C until use. For the halo assay, resultant plates were spotted with 5 μL of each protein, incubated at 37°C, and then examined for clearing zones. All experiments were performed in triplicate.

### Antibacterial activity assay

To evaluate the antibacterial activity of different LysPds against six major periodontal pathogens (*P. gingivalis* W83, *F. nucleatum* ATCC 25586, *T. denticola* ATCC 35405, *T. forsythia* ATCC 43037, *P. intermedia* ATCC 25611, and *A. actinomycetemcomitans* HK1651), each enzyme was incubated with different bacteria, respectively. Briefly, bacterial cells were grown to log phase, harvested, washed, and resuspended in buffer (10 mM HEPES, pH 7.4) to a final concentration of ∼10^6^ CFU/mL. After incubating the test bacteria with LysPds at a final concentration of 100 μg/mL for 1 h at 37°C (unless otherwise stated), viable bacteria were checked 10-fold serial dilutions onto the growth agar. All experiments were performed in quadruplicate.

Using similar culture conditions, two bacterial test strains, *P. gingivalis* W83 and *F. nucleatum* ATCC 25586, were used to evaluate the effect of different biochemical conditions on LysPds. Here, the bactericidal effect of LysPds at various concentrations (0, 6.25 μg/mL, 12.5 μg/mL, 25 μg/mL, 50 μg/mL, and 100 μg/mL), various temperatures (20 °C, 40 °C, 60 °C, 80 °C, and 100 °C) for 1 h, and stored at different temperatures (4 °C, 22 °C, or 37°C) for different time periods were tested. The antibacterial activity of LysPds after various storage times was quantified as the percentage reduction in bacterial cell count compared to the groups treated with buffer (bactericidal %), with "100%" indicating a decrease in residual cells to the detection limit. Minimum bactericidal concentration (MBC) was determined where residual cells decreased to the detection limit^68^. In addition, different pathogenic and commensal bacterial strains were used to further estimate the antimicrobial spectrum of LysPds.

### Thermophilic protein analysis

To compare the thermal stability of periodontitis candidate lysins with collected lysins, DeepTP was used to predict their thermostability^69^. Similarity, proteins from periodontal pathogens and typical clinical pathogens (listed in Extended Data Table 2) were downloaded from NCBI to compare their thermal stability.

### Circular dichroism analysis

Circular dichroism (CD) spectra of LysPds at 200 μg/mL in 10 mM HEPES (pH 7.4) were recorded on the circular dichroism spectrometer (Leatherhead, UK)^70^. Spectra were recorded over a temperature range spanning 4°C, 20°C, 40°C, 60°C, 80°C, 100°C, and 100°C-refold (heating at 100°C for 1 h followed by cooling to 4°C) covering wavelengths from 200 to 260 nm. Melting temperature (Tm) was evaluated by recording the spectra. The thermal denaturation of LysPds was assessed by monitoring the change in ellipticity from 6°C to 98°C. The CD signal was fitted by Boltzmann sigmoidal curves, which were used to calculate the Tm of LysPds^71^. All experiments were performed in triplicate.

### Transmission electron microscopy (TEM)

Log phase *P. gingivalis* W83 and *F. nucleatum* ATCC 25586 were washed and re-suspended in buffer (10 mM HEPES, pH 7.4) at a concentration of 10^8^ CFU/mL, then treated with 500 µg/mL LysPds or buffer at 37°C for 1 h. The bacterial cells were pelleted and fixed with 2.5% glutaraldehyde overnight. Finally, samples were visualized under TEM (Talos L120C, Thermo Fisher)^72^.

### Scanning electron microscopy (SEM)

To evaluate the antibacterial activity of different LysPds against planktonic bacterial cells, Log phase *P. gingivalis* W83 and *F. nucleatum* ATCC 25586 were treated as described in the TEM method. After treatment, the bacterial cells were pelleted and fixed with 2.5% glutaraldehyde overnight, visualized under SEM (SU8010, Hitachi)^72^. To investigate changes in biofilm after exposure to LysPds, biofilm models of *P. gingivalis* BNCC353909 and *F. nucleatum* ATCC 25586 in single- and dual-species were established^73^. Bacterial suspensions (10^7^ CFU/mL for *P. gingivalis*, *F. nucleatum*, and a mixture of both) were added into each 48-well plate containing cell slides, incubated at 37°C for 72 h, with media changes every 24 h. Resultant biofilm was washed with 10mM HEPES and treated with buffer or LysPds at 25 μg/mL, 100 μg/mL, and 500 μg/mL at 37°C for 1 h. The samples were visualized under SEM.

### Antibiofilm activity assay

The antibiofilm activity of LysPds was also evaluated by checking the bacterial reduction of biofilm^73^. Briefly, single-species biofilms of *P. gingivalis* BNCC353909 and *F. nucleatum* ATCC 25586 were grown and treated with buffer or LysPds at 25 μg/mL, 100 μg/mL, and 500 μg/mL at 37°C for 1 h. In addition, the wells containing the biofilm were thoroughly and vigorously pipetted to convert the biofilm cells to planktonic cells before serial dilution and plating for bacterial quantification. The antibiofilm activity of LysPds was quantified as the percentage reduction in bacterial cell count compared to the groups treated with buffer (% Clearance). "100%" indicates a decrease in residual cells to the detection limit. All experiments were performed in triplicate.

### Confocal laser scanning microscopy (CLSM)

Single- and dual-species biofilms of *P. gingivalis* BNCC353909 and *F. nucleatum* ATCC 25586 were cultivated and treated treated with buffer or LysPds at 25 μg/mL, 100 μg/mL, and 500 μg/mL at 37°C for 1h. Before microimaging, the biofilms were stained using the Live & Dead Bacterial Staining Kit^74^. The images were captured using a confocal laser scanning microscope (UltraVIEW VoX; PerkinElmer) at excitation wavelengths of 488 nm and 561 nm, with dead bacteria stained red and live bacteria stained green. All experiments were performed in triplicate.

### Metagenomic analysis of periodontitis microbiome

The study received approval from the Medical Ethics Committee of the School and Hospital of Stomatology, Wuhan University (2023LUNSHEND06). Subgingival plaque (SP) was collected from participants according to a previously established inclusion and exclusion criteria^75^. All SP samples were pooled, washed with 10mM HEPES, treated with buffer or 500 μg/mL LysPds at 37°C for 4 h, and then exposed to 100 μM propidium monoazide (PMAxx) (Biotium)^76^. DNA extraction, sequencing data processing, and species annotation were performed as previously described^77^. The downstream analysis was primarily performed and visualized using the R programming language as previously described^77^. Gram staining characteristics were predicted using Bugbase phenotypic classification^78^.

### Cytotoxicity assay

RAW 264.7 and A549 cells were cultured in Dulbecco’s modified Eagle’s medium (DMEM) supplemented with 10% fetal bovine serum (FBS) at 37°C with 5% CO_2_. The cells were seeded into 96-well culture plates at a density 10^4^ cells/well density and cultured overnight. Resultant cells were incubated with different concentrations of LysPds for 24 h, after which cell viability was assessed using a cell counting kit-8 (CCK-8). Absorbance was measured at OD_450_ using the Synergy H1 microplate reader. The medium without cells was used as a negative control for the background. Relative cell viability was determined by using the formula: %relative viability = (OD_sample_ - OD_nc_) / (OD_buffer_ - OD_nc_)] ×100%, where OD_nc_ = OD_450_ of the negative control, OD_buffer_ = OD_450_ of the buffer treated sample, and OD_sample_ = OD_450_ of the LysPds treated sample. All experiments were performed multiple times.

### Mouse infection models

The animal study was approved by the Animal Experiment Committee of the Wuhan Institute of Virology, Chinese Academy of Sciences (WIVA17202301). Mice were divided into different groups and treated according to the mouse periodontitis and calvarial infection models. Briefly, female BALB/c mice aged 6-8 weeks were fed for one week to adapt to the environment. The mice were then randomly divided into five groups, with 5 mice in each group: the normal group (Healthy), the infection group treated with 10mM HEPES (Control), the infection group treated with minocycline (Minocycline), the infection group treated with LysPd078 (LysPd078), and the infection group treated with LysPd144 (LysPd144).

The mouse periodontitis model was established following a previously described protocol^79,80^. Before the experiment, all mice were administered kanamycin in their drinking water (500 μg/mL water) for four days, followed by a 3-day antibiotic-free period. To induce periodontitis, a sterilized 5-0 silk suture was placed around the cervical margin of the right maxillary second molar. From day 0 to 13, all groups, except the normal group, were orally gavaged daily with a 100 μL mixture of *P. gingivalis* BNCC353909 and *F. nucleatum* ATCC 25586 (10^9^ CFU/mL in 2% carboxymethyl cellulose). From day 7 to day 13, mice were weighed daily, and each group was orally gavaged with either 20 μg LysPd078 (100 μL), 20 μg LysPd144 (100 μL), 10mM HEPES (100 μL), or minocycline (Periofeel Dental Ointment 2%, Showa Yakuhin Kako Co), respectively. On day 14, the mice were euthanized, and the right maxilla, heart, liver, lung, and kidney were dissected.

The mouse calvarial infection model was established following previously described^81^. To induce bone loss, a 25 μL mixture of *P. gingivalis* BNCC353909 and *F. nucleatum* ATCC 25586 (∼10^8^ CFU) was injected into the midline of the scalp between the ears of mice daily for 3 days using an insulin needle. From day 4 to day 7, each group was injected with 10 μg LysPd078 (25 μL), 10 μg LysPd144 (25 μL), 10mM HEPES (25 μL) or 1μg minocycline (25 μL). All mice were weighed daily. On day 8, mice were euthanized, and the calvarial bone, heart, liver, lung, and kidney were dissected for further analysis.

### Micro-computed tomography (micro-CT) analysis

To assess bone loss, samples were scanned with the Skyscan1276 (Bruker) at a voxel resolution of 9 μm^82^. Scanned data were reconstructed with NRecon software (Bruker). Three-dimensional images were created using CTVox software (Bruker). The distance from the cementoenamel junction (CEJ) to the alveolar bone crest (ABC) of the maxillary second mola was measured in the reconstructed 2D images with DataViewer (Bruker). The bone erosion area, including the suture calvarial area centred on the intersection of the coronal and sagittal sutures, was measured using ImageJ (NIH). Bone surface/bone volume (BS/BV) and bone volume/tissue volume (BV/TV) were quantified using CTAnalyser (Bruker).

### Histology

Maxillae and calvarial bone were fixed in 4% paraformaldehyde for 72 h and decalcified in 10% EDTA. Selected typical bones were embedded in paraffin. Immunohistochemistry staining, hematoxylin and eosin (HE) staining, tartrate-resistant acid phosphatase (TRAP) staining, and Masson’s trichrome assay were performed to evaluate the treatment effect further, following previously established methods^83,84^. Additionally, heart, liver, lung, and kidney tissues were fixed in 4% paraformaldehyde for 72 h and subjected to HE staining to assess the in vivo toxicity of treatment.

### Statistical Analyses

All calculation and statistical analyses of the experimental and computational data were conducted using GraphPad Prism V9.0 and SPSS V26.0, respectively. Statistical significance between groups was calculated using the tests indicated in each figure legend. Data are presented as mean ± standard deviation (*p < 0.05, **p < 0.01, ***p < 0.001, ****p < 0.0001, ns, not significant).

## Acknowledgments

This work was supported by the National Natural Science Foundation of China (No. 81870756), and the. We thank Dr. Yong Wang from the CAS Center for Excellence in Molecular Plant Sciences for valuable advice. We thank Dr. Xuefang An, Dr. Tao Zhand, and Dr. Li, Li at the Center for Experimental Animals, Wuhan Institute of Virology for their assistance in animal experiments. We are grateful to Dr. Ding Gao, and Dr.Bichao Xu at the Center for Instrumental Analysis and Metrology, Wuhan Institute of Virology, CAS for technical assistance in transmission electron microscopy, scanning electron microscopy, and Confocal laser scanning microscopy.

## Author information

These authors contributed equally: Fangfang Yao, Jiajun He.

## Contributions

F.Y., J.H., H.W. and Y.L. designed the study. J.H. performed the bioinformatics work.. F.Y., F.C., J.Z., carried out the experiments. F.Y., J.H., H.W. and Y.L. analyzed the data. F.Y., J.H., H.W. and Y.L. drafted the manuscript. F.Y., J.H., R.N., H.W. and Y.L. edited the manuscript.

## Corresponding authors

Correspondence to Hongping Wei and Yuhong Li

---

**Extended Data Fig. 1.**
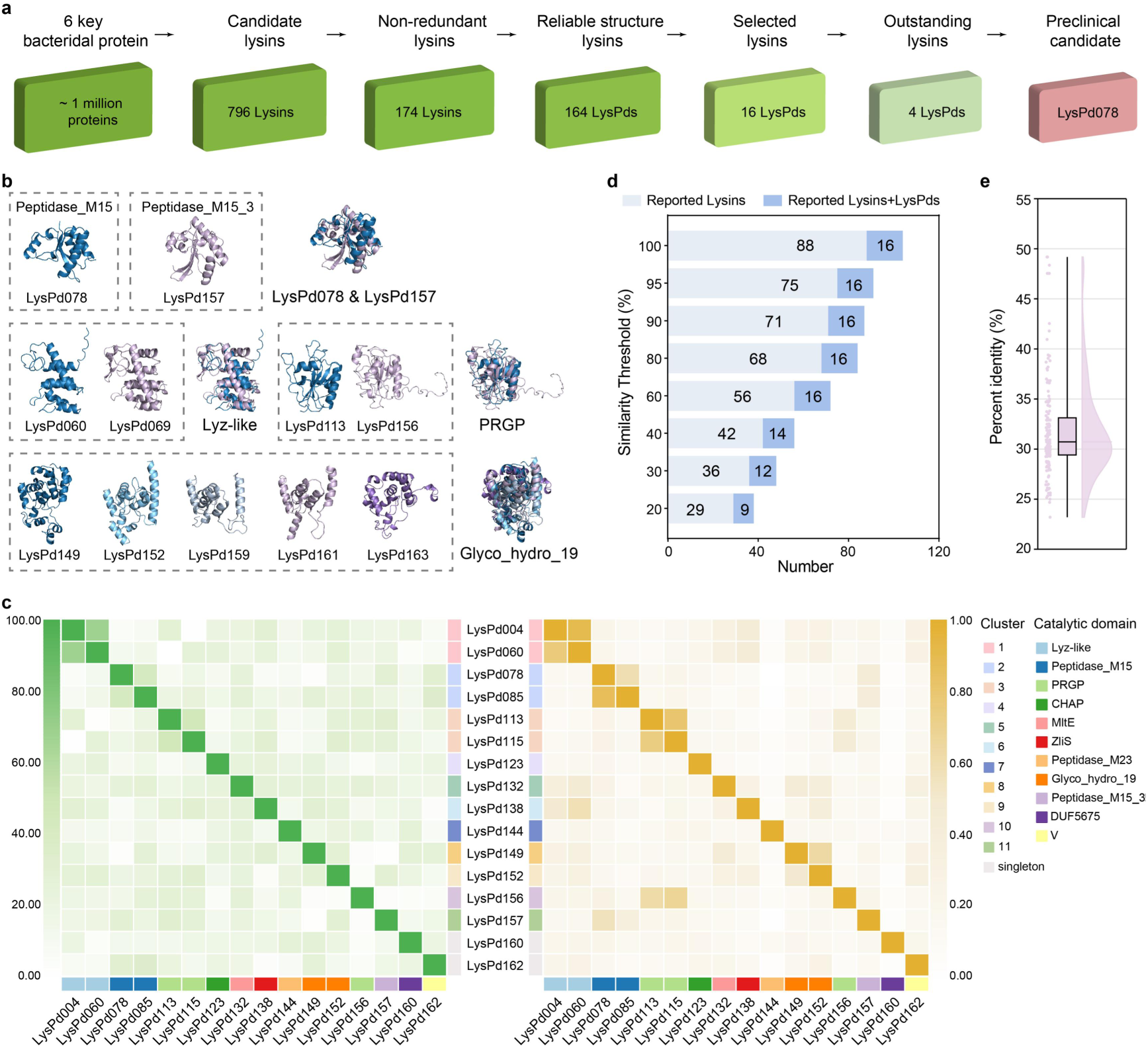
Classification of LysPds through protein structure analysis. a, Overview of the screening procedure for identifying LysPds based on structure clustering. b, Structural alignment of representative proteins from the distinct Lys_like clade (as shown in Fig. 1g) and the separate Peptidase_M15, PRGP, and Glyco_hydro_19 clusters (as shown in Fig. 2c). c, Sequence-(left) and structure-based (right) similarity matrices illustrating the similarities among the selected LysPds. The heatmap colors indicate the degree of similarity. d, Stacked bar plot showing the remaining number of selected LysPds and reported lysins after de-redundancy at different sequence similarity thresholds. e, Distribution of percent identity between 164 LysPds and reported lysins.

**Extended Data Fig. 2.**
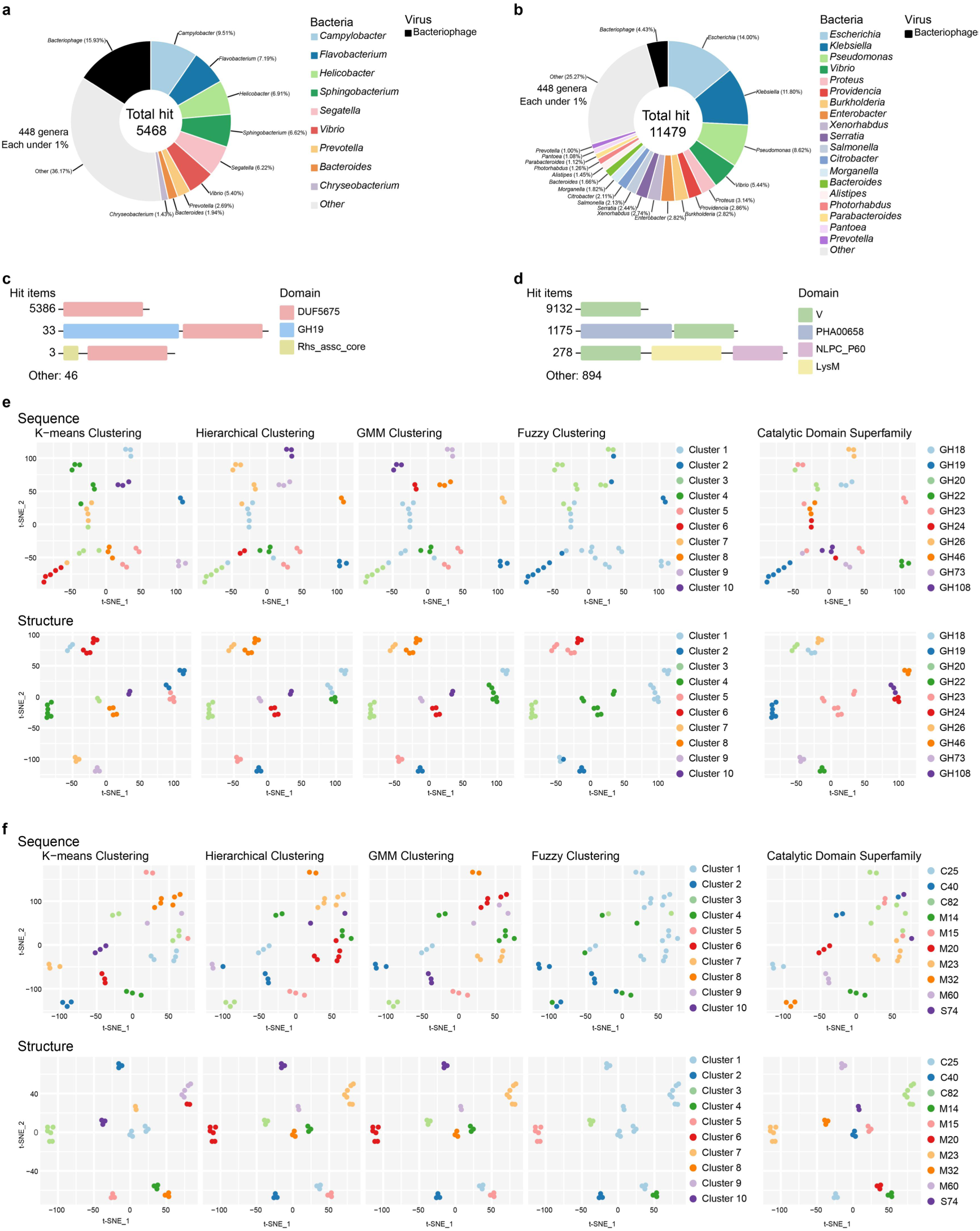
Overview of proteins containing DUF5675 domains and V domains. Distribution of DUF5675 domains (a) and V domains (b) in different bacterial genera and viruses. Schematic diagram of protein domain architecture containing DUF5675 (c) and V (d) domains, highlighting the three most abundant types. Others represent the remaining domain structure types. The sequences (Upper panels) and structures (Lower panels) of typical peptidases (e) and glycosidases (f) were clustered using various clustering methods, with a maximum of 10 cluster components.

**Extended Data Fig. 3.**
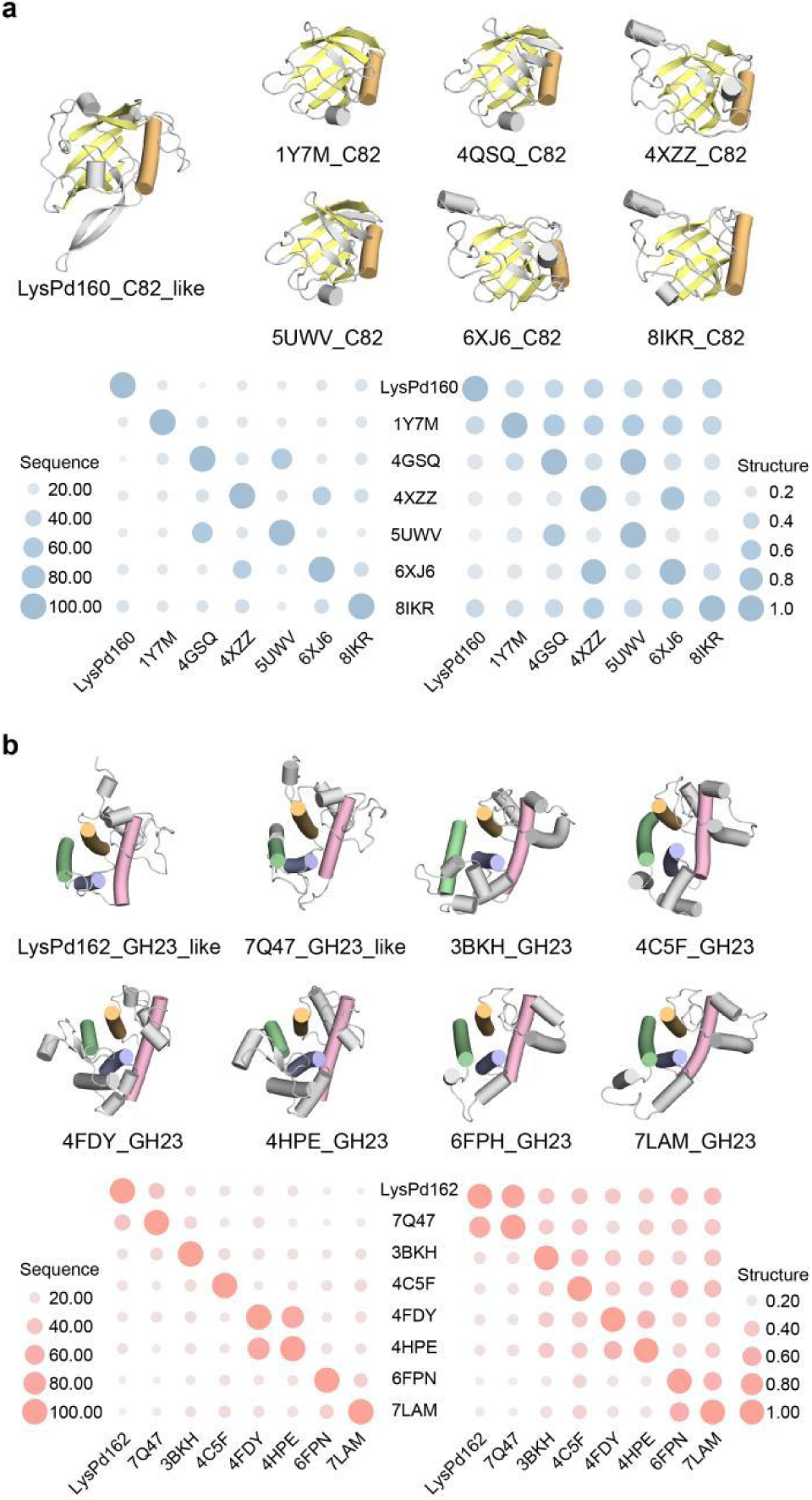
Comparison of LysPd160 with typical C82 peptidases and LysPd162 with typical GH23 glycosidases. Comparison of the core fold between LysPd160 and typical C82 peptidases (a) and between LysPd162 and typical GH23 glycosidases (b) shows conserved catalytic residues displayed as stick models, with additional protein domains removed for clarity. The sequence (left) and structural (right) similarity matrix reflects similarities between LysPd160 and typical C82 peptidases (c) and between LysPd162 and typical GH23 glycosidases (d).

**Extended Data Fig. 4.**
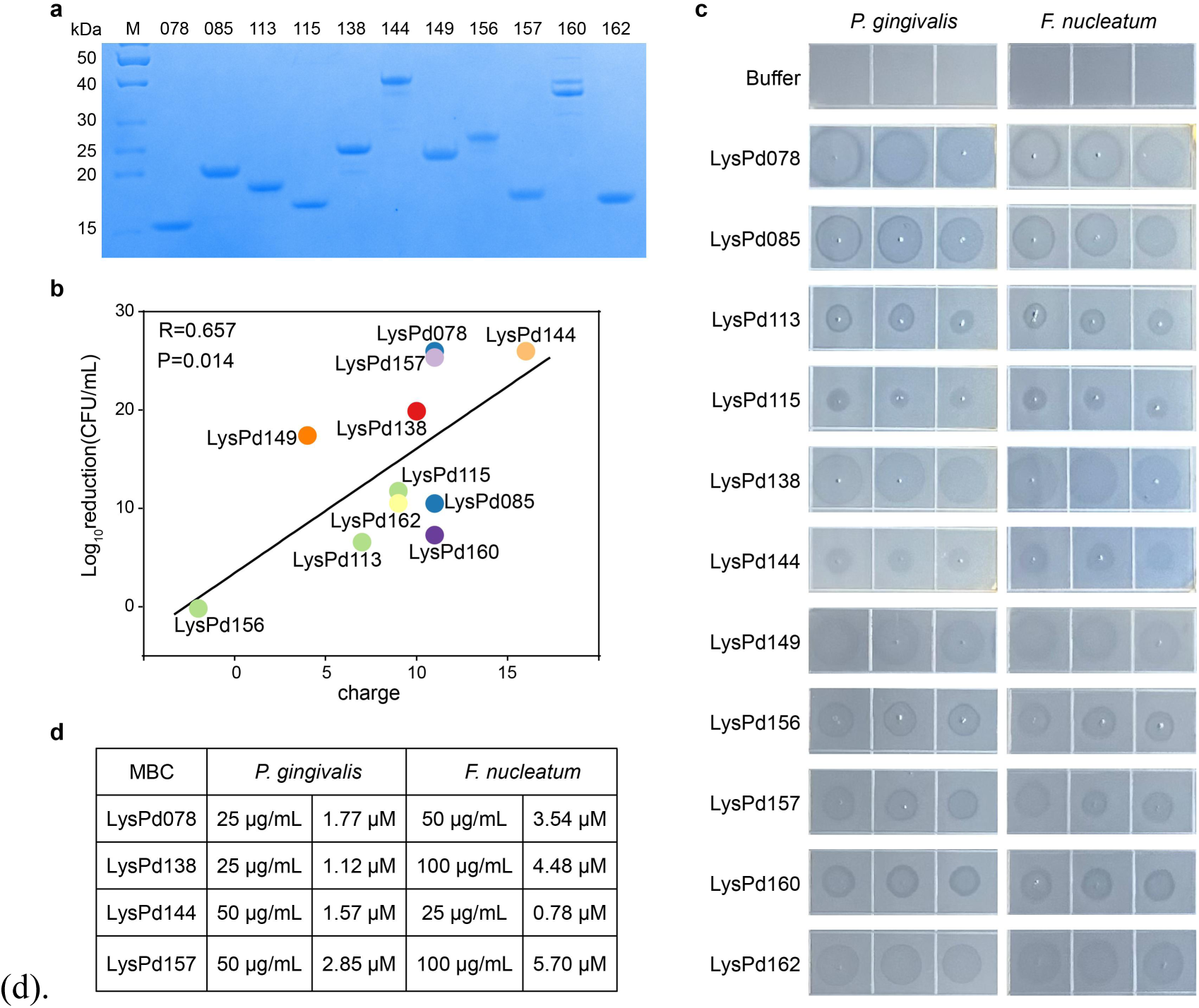
Experimental characterization of screened LysPds. a, 15% SDS-PAGE results displaying the protein bands for each LysPd selected. b, Scatter plot showing the relationship between the charge and log_10_reduction of LysPds, which corresponds to the data presented in Fig. 3a. The black line is the fitted regression line, and statistical analysis was calculated using Pearson’s correlation test (one-tailed). c, The peptidoglycan hydrolase activity of LysPds was assessed by the formation of clearing zones. d, Minimal bactericidal concentrations (MBC) were determined as the lowest concentration of lysin that resulted in the reduction of the initial inoculum to the detection limit.

**Extended Data Fig. 5.**
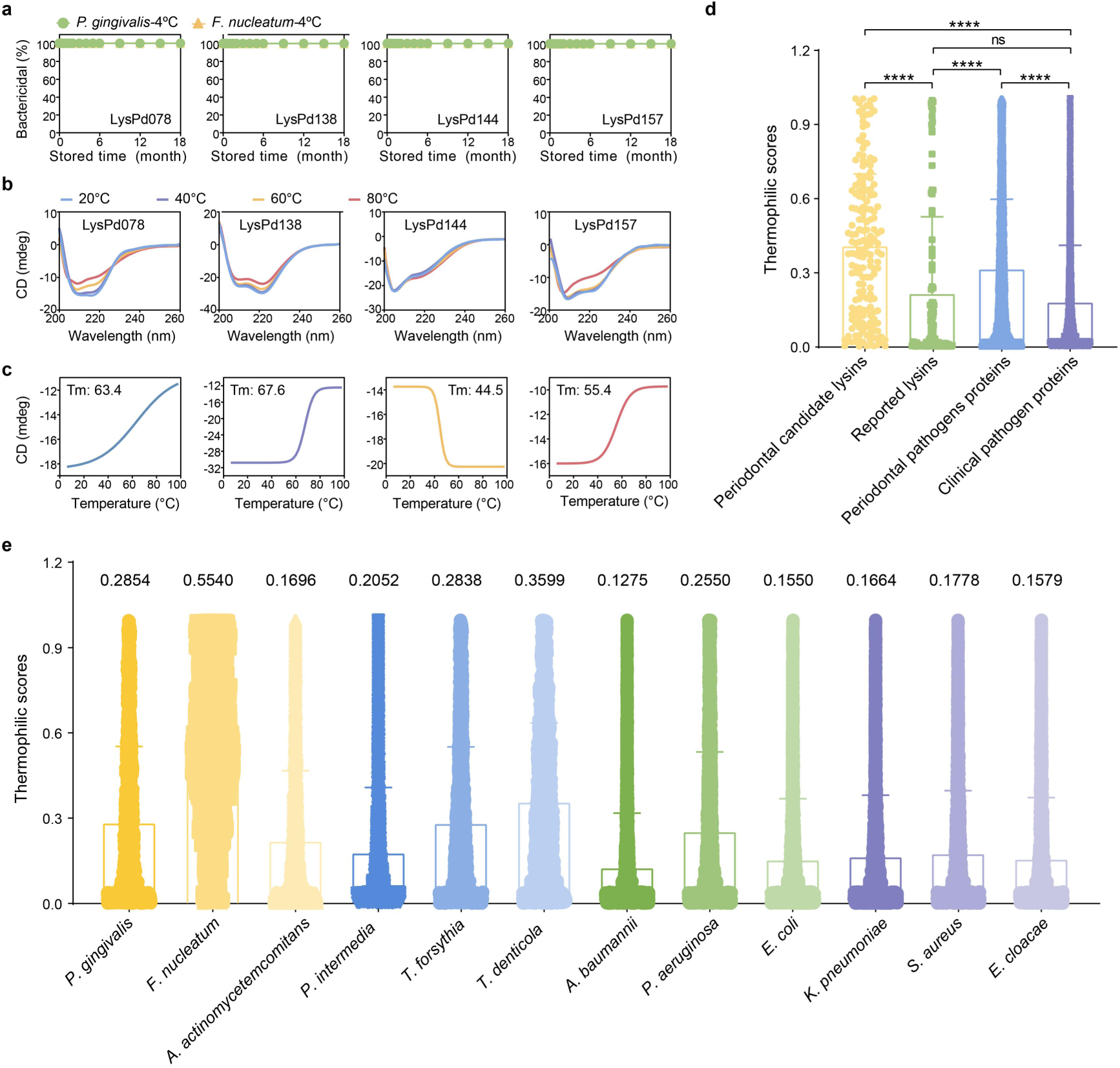
Thermal stability characterization of LysPds. a, Storage stability of LysPds at 4°C over time. b, CD spectra of LysPds at 20°C, 40°C, 60°C, and 80°C. c, Melting curve of LysPds were measured by increasing temperatures. Tm represents melting temperature, the temperature at which 50% of the protein unfolds. Statistical significance was assesed using Dunnett’s test for comparing different groups; ****p < 0.0001, ns, not significant. d, Thermophilic scores of proteins from various sources predicted by DeepTP. e, Thermophilicity of proteins from different bacterial sources. Data are presented as mean values ± SD.

**Extended Data Fig. 6.**
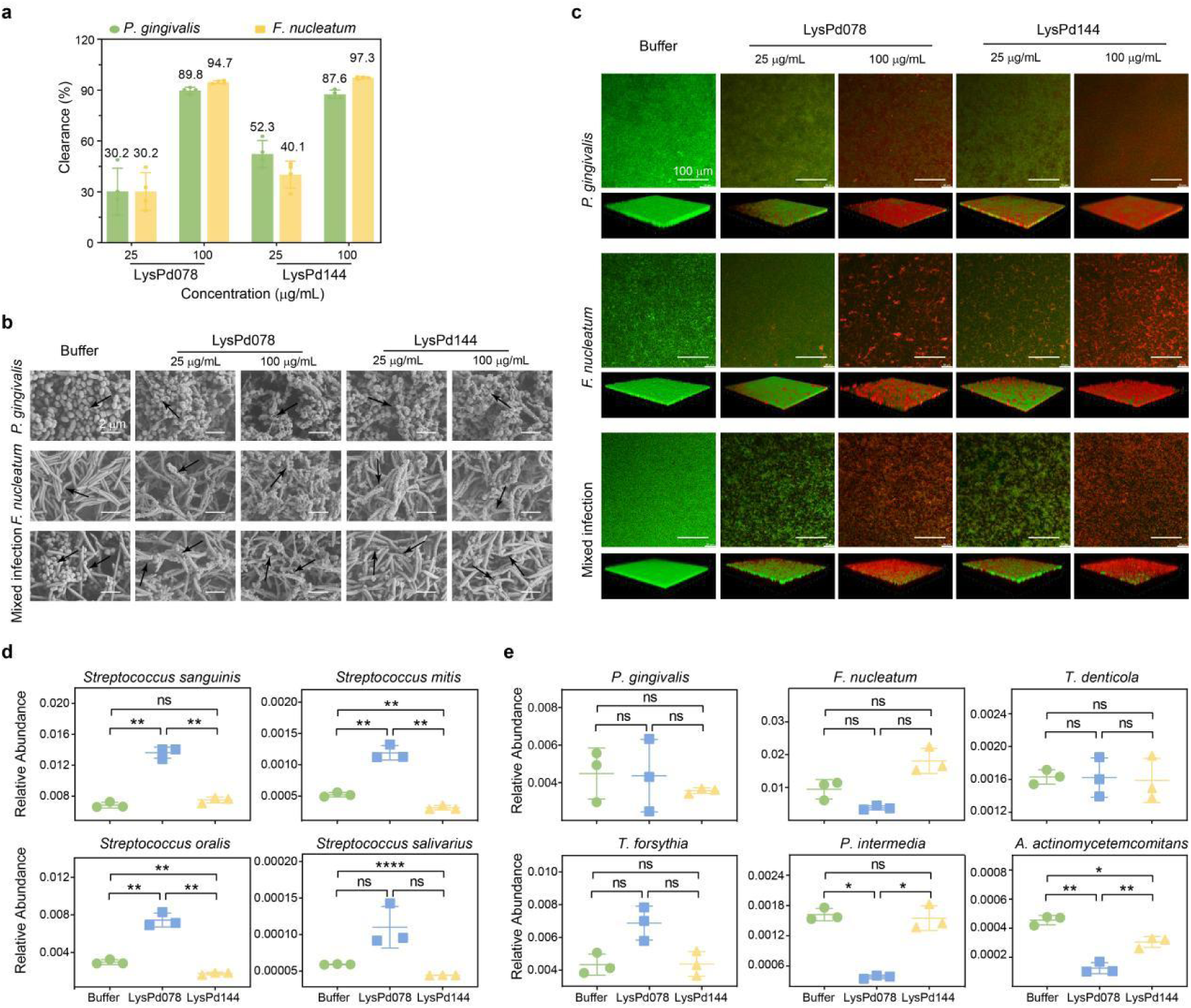
Bactericidal effect of LysPd078 and LysPd144 on biofilm models and subgingival plaque. a, Change in bacterial load within biofilms, measured as the percentage of intra-biofilm bacteria killed by LysPd078 and LysPd144, respectively. Data are presented as means ± standard deviation. b, Morphological changes of single- and dual-species biofilms treated with LysPd078 and LysPd144 visualized using SEM. Scale bar, 2 μm. c, Live/dead fluorescent staining of biofilms treated with LysPd078 and LysPd144. Green fluorescence indicated live bacteria, and red fluorescence indicated dead bacteria. Scale bar, 100 μm. Experiments were performed in duplicates with similar results, and one representative figure is shown. The relative abundances of different groups at the species level in typical oral commensal bacteria (d) and periodontitis pathogenic bacteria (e). Each data point represents a biologically independent sample. Statistical significance was calculated using Permutation test; n = 3/group, *p < 0.05, **p < 0.01, ns, not significant.

**Extended Data Fig. 7.**
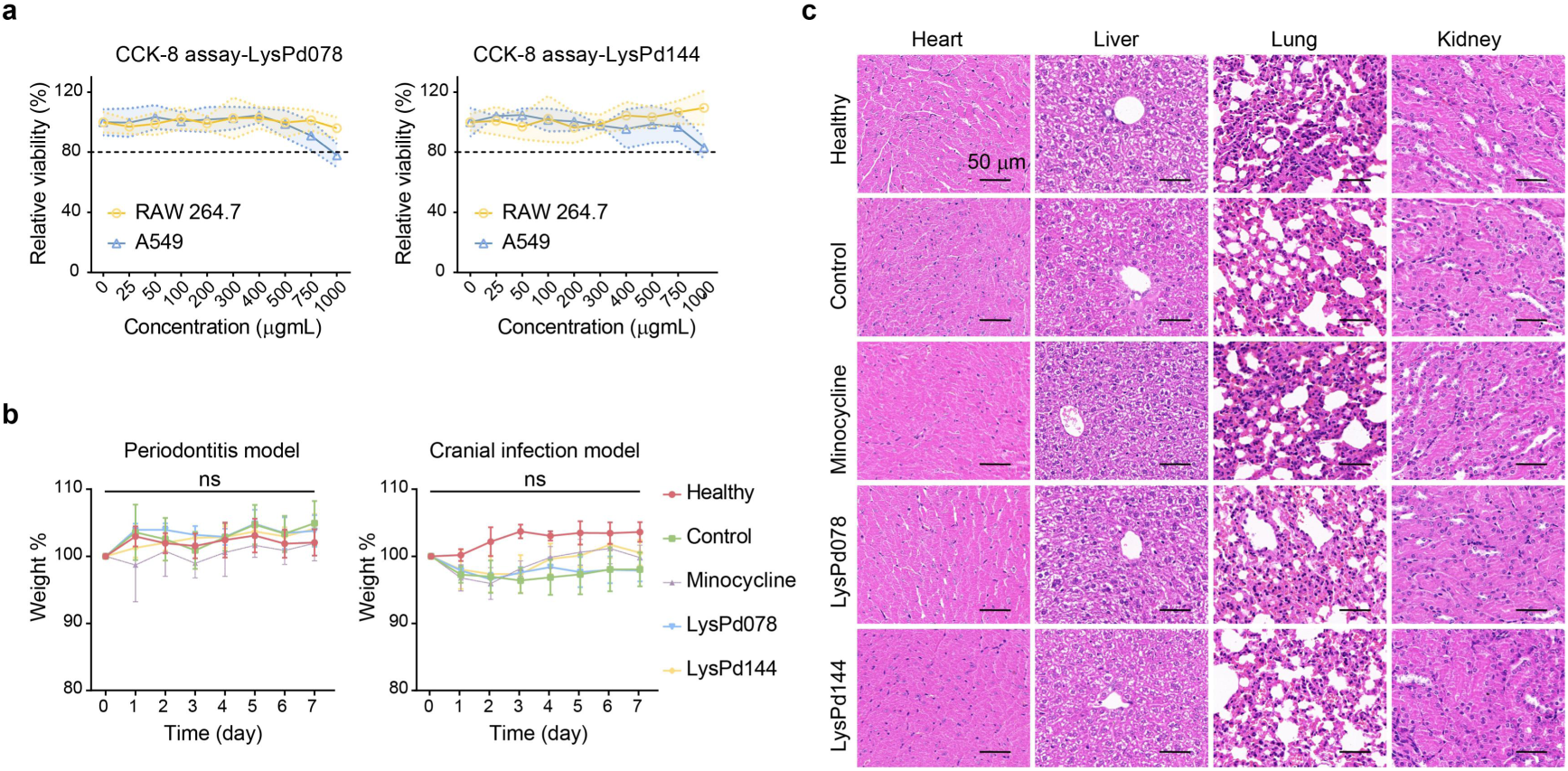
In vitro and in vivo safety assessment of LysPd078 and LysPd144. a, Relative cell viability of RAW264.7 cells and A549 cells compared to the buffer-treated control. Lines and shaded regions indicate means and SD, respectively. b, Changes in body weight among Healthy mice and those treated with different conditions. Statistical significance was calculated according to One Way ANOVA; n = 5/group, ns, not significant. Data are presented as mean values ± SD. c, Histological evaluation of representative organs (heart, liver, lung, kidney) stained with H&E in periodontitis model mice. Scale bar, 50 µm.

**Extended Data Table 1.** Characteristics of Thermostable lysins.

| Lysin | Origin | Observations<br>(temperature, time, remain activity) |
| --- | --- | --- |
| LysPd078 | *Fusobacterium nucleatum* prophage | 100°C, 1 h, ~100% activity |
| LysPd138 | *Porphyromonas gingivalis* prophage | 100°C, 1 h, ~100% activity |
| LysPd144 | *Porphyromonas gingivalis* prophage | 100°C, 1 h, ~100% activity |
| LysPd157 | *Prevotella intermedia* prophage | 100°C, 1 h, ~100% activity |
| **Thermophilic phages** |  |  |
| TP84_28<sup>85</sup> | *Geobacillus stearothermophilus* phage TP-84 | 100°C, 0.5 h, trace activity |
| GVE2<sup>86</sup> | Thermophilic phage GVE2 | 60°C, 1 h, ~40% activity |
| TSPphg<sup>87</sup> | *Thermus* phage TSP4 | 85°C, 0.5 h, ~15% activity |
| PhiKo<sup>88</sup> | *Thermus thermophilus* phage φKo | 100°C, 5 h, > 77% activity |
| Ts2631<sup>89</sup> | *Thermus scotoductus* phage vB_Tsc2631 | 95°C, 2 h, > 64.8% activity |
| Ph2119<sup>90</sup> | *Thermus scotoductus* phage Ph2119 | 95°C, 6 h, > 87% activity |
| MMPphg<sup>91</sup> | *Thermophilic Meiothermus* phage MMP17 | 70°C, 0.5 h, ~50% activity |
| φIN93<sup>92</sup> | *Thermus aquaticus* TZ2 phage φIN93 | 95°C, 1 h, ~38.6% activity |
| AmiP<sup>93</sup> | *Hyperthermophilic Thermus parvatiensis* prophage | 99°C, 0.5 h, ~92.9% activity |
| **Other phage** |  |  |
| gp36C<sup>94</sup> | *Pseudomonas aeruginosa* phage φKMV | 100°C, 1 h, ~50% activity |
| LysWL59<sup>95</sup> | *Salmonella Typhimurium* phage LPST10 | 90°C, 1 h, ~75% activity |
| Gp110<sup>66</sup> | *Salmonella* phage 10 | 90°C, 1 h, ~26.7% activity |
| KP27<sup>96</sup> | *Klebsiella pneumoniae* phages vB_KpnM_KP15 | 90°C, 1 h, ~50% activity |
| Lys68<sup>71</sup> | *Salmonella* phage phi68 | 100°C, 0.5 h, ~10.2% activity |
| PVP-SE1<sup>97</sup> | *Salmonella enterica* phage PVP-SE1 | 90°C, 1 h, ~10% activity |

**Extended Data Table 2.**
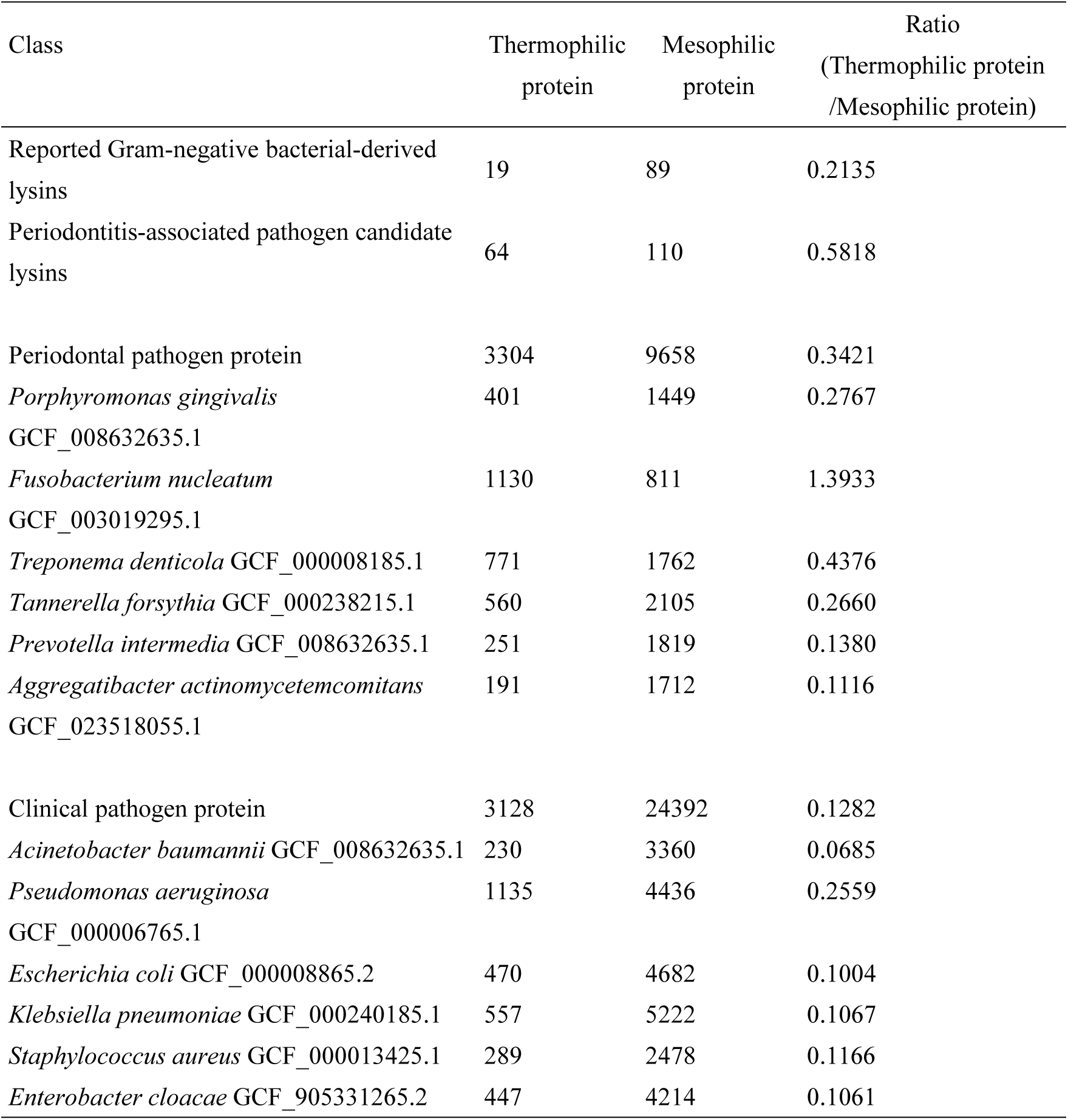
Thermophilic scores of different bacterial groups predicted by DeepTP.

