## Supplementary Tables for "Structure-based similarity network accelerates the discovery of lysins as oral microbiome modulators targeting periodontal pathogens"

**Supplementary Table 1. Bacterial strains and growth conditions.**

| Species | Strain | Medium[a] | Condition[b] | Source[c] |
|---|---|---|---|---|
| Oral pathogens | | | | |
| *Aggregatibacter actinomycetemcomitans* | BNCC337071 | 2 | 3 | 2 |
| | HK1651 | 2 | 3 | 2 |
| *Enterococcus faecalis* | ATCC 29212 | 1 | 3 | 2 |
| | ATCC 51299 | 1 | 3 | 2 |
| | WHS30008 | 1 | 3 | 1 |
| *Fusobacterium nucleatum* | ATCC 10953 | 2 | 4 | 2 |
| | ATCC 25586 | 2 | 4 | 2 |
| *Porphyromonas gingivalis* | ATCC 33277 | 2 | 4 | 2 |
| | BNCC353909 | 2 | 4 | 2 |
| | W83 | 2 | 4 | 2 |
| *Prevotella intermedia* | ATCC 25611 | 2 | 4 | 2 |
| | BNCC352061 | 2 | 4 | 2 |
| *Streptococcus mutans* | UA159 | 1 | 3 | 2 |
| *Treponema denticola* | ATCC 35405 | 3 | 4 | 3 |
| *Tannerella forsythia* | ATCC 43037 | 4 | 4 | 4 |
| Oral commensals | | | | |
| *Streptococcus mitis* | 59 | 1 | 3 | 2 |
| *Streptococcus oralis* | BNCC354691 | 1 | 3 | 2 |
| | 6715 | 1 | 3 | 2 |
| | WHS21020 | 1 | 3 | 1 |
| | WHS21021 | 1 | 3 | 1 |
| *Streptococcus sanguis* | 641 | 1 | 3 | 2 |
| | ATCC 10556 | 1 | 3 | 2 |
| Other pathogens | | | | |
| *Acinetobacter baumannii* | WHG40035 | 5 | 1 | 1 |
| | WHG40137 | 5 | 1 | 1 |
| *Bacillus cereus* | WHG21001 | 5 | 1 | 1 |
| *Citrobacter amalonaticus* | WHG10063 | 5 | 1 | 1 |
| *Dickeya dadantii* | WHG10105 | 5 | 2 | 1 |
| *Dickeya fangzhongdai* | WHG10102 | 5 | 2 | 1 |
| | WHG10103 | 5 | 2 | 1 |
| *Enterobacter cloacae* | WHG10064 | 5 | 1 | 1 |
| *Enterococcus faecium* | WHS30003 | 1 | 1 | 1 |
| | WHS30005 | 1 | 1 | 1 |
| *Escherichia coli* | WHG10002 | 5 | 1 | 1 |
| | WHG10003 | 5 | 1 | 1 |
| *Helicobacter pylori* | SS1 | 6 | 3 | 1 |
| *Klebsiella pneumoniae* | WHG11004 | 5 | 1 | 1 |
| | WHG11009 | 5 | 1 | 1 |
| *Morganella morganii* | WHG10062 | 6 | 1 | 1 |

| | | | | |
|---|---|---|---|---|
| *Pectobacterium aroidearum* | WHG10100 | 5 | 2 | 1 |
| | WHG10101 | 5 | 2 | 1 |
| *Pectobacterium polaris* | WHG10104 | 5 | 2 | 1 |
| *Proteus mirabilis* | WHG10061 | 7 | 1 | 1 |
| *Pseudomonas aeruginosa* | WHG50021 | 5 | 1 | 1 |
| | WHG50035 | 5 | 1 | 1 |
| *Pseudomonas koreensis* | WHG10106 | 5 | 2 | 1 |
| *Ralstonia solanacearum* | GIM1.74 | 8 | 2 | 1 |
| *Salmonella enterica* | ATCC 13076 | 5 | 1 | 1 |
| | CICC (B) 21497 | 5 | 1 | 1 |
| *Salmonella paratyphi* | CMCC (B) 50094 | 5 | 1 | 1 |
| *Salmonella Typhimurium* | CMCC (B) 50115 | 5 | 1 | 1 |
| *Shigella flexneri* | M90T | 5 | 1 | 1 |
| *Shigella sonnei* | WHG10065 | 5 | 1 | 1 |
| *Staphylococcus aureus* | N315 | 1 | 1 | 1 |
| | T23 | 1 | 1 | 1 |
| *Staphylococcus epidermidis* | WHS10011 | 1 | 1 | 1 |
| *Staphylococcus nepalensis* | WHS11098 | 1 | 1 | 1 |
| *Streptococcus agalactiae* | S12 | 1 | 3 | 1 |
| *Streptococcus pneumoniae* | WHS20011 | 1 | 3 | 1 |
| | WHS20013 | 1 | 3 | 1 |
| *Streptococcus pyogenes* | ATCC 49399 | 1 | 3 | 1 |
| *Streptococcus suis* | WHS21001 | 1 | 3 | 1 |
| *Vibrio parahaemolyticus* | ATCC 17802 | 1 | 1 | 1 |
| *Vibrio vulnificus* | ATCC 27562 | 1 | 1 | 1 |
| Other commensals | | | | |
| *Bifidobacterium animalis subsp. lactis* | WHG91006 | 9 | 4 | 1 |
| *Lacticaseibacillus rhamnosus* | WHG91001 | 9 | 3 | 1 |
| *Lactiplantibacillus plantarum* | WHG91002 | 9 | 3 | 1 |
| *Lactobacillus delbrueckii subsp. bulgaricus* | WHG91005 | 9 | 3 | 1 |
| *Ligilactobacillus salivarius salivarius* | WHG91004 | 9 | 3 | 1 |
| *Loigolactobacillus coryniformis* | WHG91003 | 9 | 3 | 1 |
| *Streptococcus thermophilus* | WHS21010 | 9 | 3 | 1 |
| Other | | | | |
| *Escherichia coli* | BL21 (DE3) | 5 | 1 | 1 |

[a]Media: 1. Tryptic Soy Broth. 2.Tryptic Soy Broth supplemented with 5 mg/mL yeast extract, 0.5 mg/mL L-cysteine hydrochloride, 5 μg/mL hemin and 1 μg/mL menadione. 3. Fastidious Anaerobe Broth (Solarbio, China). 4. Fastidious Anaerobe Broth supplemented with 1 μg/mL MurNAc. 5. Lysogeny broth (LB). 6. Tryptic Soy Broth with 5% FBS. 7. Lysogeny broth (LB) with 0.1% borate. 8. Casamino acid-Peptone-Glucose broth. 9. de Man, Rogosa and Sharpe Broth. The agar plate of media 2, 3, 4, 6 were supplemented with 5% defibrinated sheep blood.

[b]Condition: 1. Aerobic cultivation, 37°C with aeration (200 rpm). 2. Aerobic cultivation, 28°C with aeration (200 rpm). 3. Microaerophilic cultivation, 37°C under 5% $CO_2$, 4. Anaerobic cultivation, 37°C with AnaeroPack™ (Mitsubishi, Japan).

[c]Source: 1. Key Laboratory of Special Pathogens and Biosafety, Wuhan Institute of Virology, China. 2. Stomatological Hospital of Wuhan University, China. 3. Oral Microbiome Bank of China. 4. Shanghai Ninth People's Hospital of Shanghai Jiao Tong University, China.

**Supplementary Table 2. Reported natural lysins against Gram-negative bacteria.**

| Reported lysin | Origin | Catalytic domain | Lenth (aa) |
|---|---|---|---|
| 201phi2-1gp229 | *Pseudomonas aeruginosa* phage 201phi2-1 | Muramidase | 262 |
| AP3gp15 | *Burkholderia* phage AP3 | Muramidase | 266 |
| BcepC6Bgp22 | *Burkholderia cepacia* phage BcepC6B | COG4678 | 164 |
| Bp7e | *Escherichia coli* phage Bp7 | Lyz-like | 162 |
| BSP16Lys | *Salmonella* phage BSP161 | PGRP | 151 |
| EL188 | *Pseudomonas aeruginosa* phage φ EL | Lyz-like | 292 |
| ElyA1 | *Acinetobacter baumannii* phage Ab105 φ1 | ZliS | 181 |
| EndoRB43 | *Escherichia coli* phage RB43 | Peptidase_M15 | 131 |
| EndoRB49 | *Escherichia coli* phage RB49 | Peptidase_M15 | 129 |
| Gp110 | *Salmonella* phage 10 | Muramidase | 264 |
| gp61 | *Escherichia coli* phage N4 | ZliS | 208 |
| K11gp3.5 | *Klebsiella pneumoniae* K11 | PGRP | 151 |
| KP27 | *Klebsiella pneumoniae phages* vB_KpnM_KP15 | Peptidase_M15 | 131 |
| KP32gp15 | *Klebsiella pneumoniae* phage KP32 | PGRP | 151 |
| KZ144 | *Pseudomonas aeruginosa* phage φKZ | PGRP | 260 |
| LyS15S6 | *Salmonella* phage BPS15S6 | Lyz-like | 154 |
| Lys394 | *Salmonella* phage S-394 | Peptidase_M15 | 137 |
| Lys68 | *Salmonella* phage Φ68 | Lyz-like | 162 |
| LysB4 | *Bacillus cereus* phage B4 | Peptidase_M15 | 262 |
| LysCs4 | *Cronobacter sakazakii* strain ATCC BAA-894 | Lyz-like | 164 |
| LysECP26 | *Escherichia coli* phage vB_EcoM-ECP26 | Lyz-like | 156 |
| Lysep3 | *Escherichia coli* phage vB_EcoM-ep3 | Lyz-like | 159 |
| LysF1 | *Escherichia coli* phage FAHEc1 | Lyz-like | 154 |
| LysO78 | *Escherichia coli* vB_EcoM_APEC | Glyco_hydro_19 | 202 |
| LysPN09 | *Pseudomonas syringae* phage PN09 | Muramidase | 185 |
| Lysqdvp001 | *Vibrio parahaemolyticus* phage qdvp001 | NLPC_P60 | 236 |
| LysSE24 | *Salmonella* phage SLMP1 | Lyz-like | 162 |
| LysSs1 | *Cronobacter* phage vB_CsaP_Ss1 | Peptidase_M15 | 135 |
| LysSTP4 | *Salmonella enterica* phage STP4-a | Lyz-like | 166 |

| | | | |
|---|---|---|---|
| LysVpKK5 | *Vibrio parahaemolyticus* phage VpKK5 | PGRP | 163 |
| LysWL60 | *Salmonella Typhimurium* phage LPST10 | Lyz-like | 92 |
| M15A | *Burkholderia pseudomallei* phage ST79 | Peptidase_M15_3 | 149 |
| M4Lys | *Salmonella Typhimurium* phage BSPM4 | V | 237 |
| p02 | *Pseudomonas aeruginosa* phage PaP3 | Lyz-like | 165 |
| P2gp09 | *Escherichia coli* phage P2 | Lyz-like | 165 |
| PaP1 | *Pseudomonas aeruginosa* phage PaP1 | Muramidase | 186 |
| Ph2119 | *Thermus scotoductus* phage Ph2119 | Amidase_2 | 155 |
| Ply17 | *Pseudomonas aeruginosa* phage φYY | Lyz-like | 230 |
| PlyM22 | Viral Metagenome | Lyz-like | 159 |
| PlyPAJD-1 | *Pseudomonas aeruginosa* phage vB_PaeS-PAJD-1 | Lyz-like | 174 |
| PsP3gp10 | *Salmonella enterica* phage PsP3 | Lyz-like | 165 |
| PVP-SE1gp | *Salmonella enterica* phage PVP-SE1 | Glyco_hydro_19 | 184 |
| SPN1S | *Salmonella enterica* phage SPN1S | Glyco_hydro_19 | 209 |
| Spp62 | *Shewanella putrefaciens* Phage Spp001 | Hydrolase_2 | 208 |
| T5 | *Escherichia coli* phage T5 | Peptidase_M15 | 137 |
| T7 | *Escherichia coli* phage T7 | PGRP | 151 |
| ABgp46 | *Acinetobacter baumannii* phage vb_AbaP_CEB1 | Glyco_hydro_19 | 185 |
| Abtn-4 | *Acinetobacter baumannii* phage vB_AbaP_D2 | Glyco_hydro_19 | 185 |
| AcLys | *Acinetobacter baumannii* AB 5075 strain prophage | Lyz-like | 184 |
| CfP1 | *Citrobacter freundii* phage CfP1 | Peptidase_M15 | 131 |
| LysAB2 | *Acinetobacter baumannii* phage φAB2 | Glyco_hydro_19 | 185 |
| LysAB3 | *Acinetobacter baumannii* ATCC 17978 strain prophage | Lyz-like | 182 |
| LysAB4 | *Acinetobacter baumannii* ATCC 17978 strain prophage | Lyz-like | 169 |
| LysABP-01 | *Acinetobacter baumannii* phage φABP-01 | Glyco_hydro_19 | 185 |
| LysAm24 | *Acinetobacter baumannii* phage AM24 | Lyz-like | 224 |
| LysAp22 | *Acinetobacter baumannii* phage AP22 | Lyz-like | 184 |
| LysECD7 | *Escherichia coli* phage ECD7 | Peptidase_M15 | 129 |
| LysMK34 | *Acinetobacter baumannii* phage MK34 | Glyco_hydro_19 | 185 |
| LysPA26 | *Pseudomonas aeruginosa* phage JD010 | Lyz-like | 145 |
| LysSAP26 | *Staphylococcus aureus* phage SAP-26 | CHAP | 251 |
| LysSi3 | *Salmonella enterica* phage UAB_Phi87 | Lyz-like | 154 |

| | | | |
|---|---|---|---|
| LysSP1 | *Salmonella Typhimurium* phage SLMP1 | Lyz-like | 162 |
| LysSS | *Salmonella enterica* phage SS3e | Lyz-like | 162 |
| LysSTG2 | *Salmonella Typhimurium* phage STG2 | Peptidase_M15 | 137 |
| LysWL59 | *Salmonella Typhimurium* phage LPST10 | Lyz-like | 154 |
| Mitrecin A | *Streptomyces* sp. 212 strain prophage | Peptidase_M15 | 127 |
| MMPphg | *Thermophilic Meiothermus* phage MMP17 | NlpD | 210 |
| OBPgp279 | *Pseudomonas fluorescens* phage OBP | Glyco_hydro_19 | 328 |
| P28 | *Stenotrophomonas maltophilia* phage P28 | Lyz-like | 161 |
| Pae87 | *Pseudomonas aeruginosa* phage JG004 | Muramidase | 186 |
| Ply6A3 | *Acinetobacter baumannii* phage vB_AbaP_PD-6A3 | Glyco_hydro_19 | 185 |
| PlyAB1 | *Acinetobacter baumannii* phage Abp1 | Glyco_hydro_19 | 185 |
| PlyE146 | *Escherichia coli* 8.0569 strain prophage | Lyz-like | 146 |
| PlyEc2 | *Escherichia coli* phage | Glyco_hydro_19 | 209 |
| PlyF307 | *Acinetobacter baumannii* phage RL-2015 | Lyz-like | 146 |
| PlyPa01 | *Pseudomonas aeruginosa* strain prophage | Lyz-like | 143 |
| PlyPa03 | *Pseudomonas aeruginosa* strain prophage | RrrD | 144 |
| PlyPa91 | *Pseudomonas aeruginosa* strain prophage | Lyz-like | 154 |
| PlyPa96 | *Pseudomonas aeruginosa* strain prophage | RrrD | 170 |
| pPHAb1 | *Acinetobacter baumannii* strain prophage | Muramidase | 276 |
| pPHAb2 | *Acinetobacter baumannii* strain prophage | Muramidase | 284 |
| pPHAb3 | *Acinetobacter baumannii* strain prophage | Muramidase | 276 |
| pPHAb4 | *Acinetobacter baumannii* strain prophage | Muramidase | 286 |
| pPHAb5 | *Acinetobacter baumannii* strain prophage | Muramidase | 285 |
| pPHAb6 | *Acinetobacter baumannii* strain prophage | Muramidase | 196 |
| SPN9CC | *Salmonella Typhimurium* phage SPN9CC | Lyz-like | 167 |
| Ts2631 | *Thermus scotoductus* phage vB_Tsc2631 | Amidase_2 | 156 |
| TSPphg | Thermus phage TSP4 | NlpD | 166 |

**Supplementary Table 3. The most matching sequence of reported lysins for selected LysPds.**

| Cluster | Lysin | Query Len | Matching Reported lysin | Hit Len | Cover % | E-value | Identity% |
|---|---|---|---|---|---|---|---|
| 1 | LysPd004 | 171 | AcLys | 184 | 87 | 7.31E-09 | 27.33 |
| 1 | LysPd060 | 146 | LysAp22 | 184 | 92 | 1.40E-07 | 28.67 |
| 2 | LysPd078 | 122 | EndoRB49 | 129 | 100 | 1.43E-42 | 48.36 |
| 2 | LysPd085 | 169 | EndoRB49 | 129 | 95 | 2.03E-19 | 31.88 |
| 3 | LysPd113 | 157 | LysVpKK5 | 163 | 93 | 3.84E-21 | 31.29 |
| 3 | LysPd115 | 147 | KP32gp15 | 151 | 95 | 2.32E-25 | 33.80 |
| 4 | LysPd123 | 138 | LysSAP26 | 251 | 96 | 8.77E-12 | 31.47 |
| 5 | LysPd132 | 190 | Ply17 | 230 | 49 | 4.62E-08 | 32.14 |
| 6 | LysPd138 | 191 | ElyA1 | 181 | 63 | 8.39E-07 | 29.63 |
| 7 | LysPd144 | 284 | MMPphg | 210 | 33 | 7.96E-09 | 33.33 |
| 8 | LysPd149 | 193 | PVP-SE1gp | 184 | 97 | 2.26E-24 | 34.34 |
| 9 | LysPd152 | 159 | LysABP-01 | 185 | 100 | 2.62E-23 | 38.82 |
| 10 | LysPd156 | 208 | BSP16Lys | 151 | 55 | 0.008 | 25.41 |
| 11 | LysPd157 | 153 | M15A | 149 | 80 | 8.72E-24 | 40.95 |
| singleton | LysPd160 | 294 | EL188 | 292 | 37 | 3.64E-10 | 36.61 |
| singleton | LysPd162 | 137 | M4Lys | 237 | 83 | 1.47E-17 | 35.77 |
